# Diencephalic integrity explains aspects of hippocampal amnesia

**DOI:** 10.1101/2025.05.21.655297

**Authors:** Georgios P. D. Argyropoulos, John P. Aggleton, Christopher R. Butler

**Affiliations:** Memory Research Group, Nuffield Department of Clinical Neurosciences, University of Oxford, OX3 9DU, UK; Division of Psychology, Faculty of Natural Sciences, University of Stirling, Scotland, FK9 4LA, UK; School of Psychology, Cardiff University, CF10 3AT, UK; The George Institute for Global Health, Imperial College London, W12 7RZ, UK; Department of Brain Sciences, Imperial College London, W12 0NN, UK; Departamento de Neurología, Pontificia Universidad Católica de Chile, Chile

**Keywords:** amnesia, hippocampus, mammillary bodies, thalamus, white matter

## Abstract

Research on amnesia has been fundamental in establishing the role of the human hippocampus in memory. Even though other structures within the hippocampal-diencephalic-cingulate network also play a role in episodic memory, studies of hippocampal amnesia ignore the importance of damage in this broader network. In a large cohort of patients (n=38) with hippocampal damage due to autoimmune limbic encephalitis, we had found the amnesia was predominantly explained by abnormalities in this network. Here, we examined the integrity of specific diencephalic nuclei and of white matter pathways, and whether it explains patients’ amnesia. We found atrophy in the anterior thalamic nuclei and the mammillary bodies, but also in the laterodorsal, pulvinar, and dorsomedial nuclei. Atrophy was often as pronounced as that in the hippocampal formation, even though these patients are often studied as “human models” of hippocampal damage. Diencephalic volumes predicted memory over and above any hippocampal/subicular subfield volume estimate. White matter was compromised within and beyond this network. Fornix integrity was linked to diencephalic and hippocampal volumes, but not to recollection/recall. We strongly advise caution in employing the term “focal hippocampal damage”, and highlight the need to study the significance of plausibly knock-on effects in diencephalic nuclei within broader circuits.

## Introduction

Hippocampal amnesia has taken centre stage in memory neuroscience since patient H.M (Scoville and Milner 1957), and is commonly explained on the basis of hippocampal volume reduction (Wixted and Squire 2004; Gold et al. 2006; Patai et al. 2015; Dede et al. 2016; Miller et al. 2017). However, recent work (echoing earlier insight: Warrington and Weiskrantz, 1982; Heilman et al., 1990; Clarke et al., 1994) emphasises functional dysconnectivity between the affected hippocampal formation and the rest of the brain as explaining hippocampal amnesia (Hayes et al. 2012; Rudebeck et al. 2013; Henson et al. 2016; Miller et al. 2020). Moreover, there is evidence for abnormalities beyond the medial temporal lobe that may follow hippocampal damage (Cormack et al. 2005; Rosenberg et al. 2006; Mueller et al. 2010; Morgan et al. 2015; Romeo et al. 2019; Tung et al. 2022). Extra-hippocampal damage within the “extended hippocampal system” (Aggleton and Brown, 1999; “hippocampal-diencephalic-cingulate network”: Bubb et al., 2017; “revised Papez circuit”: Aggleton et al., 2022), may also impair episodic memory (Aggleton and Brown 1999; Tsivilis et al. 2008; Kafkas et al. 2020): fornical damage may impair recollection (retrieval of contextual details of events) and thus recall (Carlesimo et al. 2007; Tsivilis et al. 2008; Vann et al. 2009), but not familiarity (the sense of having encountered a stimulus, without retrieving contextual details) and recognition (supported by recollection and familiarity).

Despite acknowledging the need to examine this network across memory-impaired patients (Jonin et al. 2018; Basile et al. 2020), there remains very little research on the explanatory potential of such knock-on effects in hippocampal amnesia (Argyropoulos et al. 2019; Harms et al. 2023). This perspective may help resolve debates on hippocampal contributions to processing remote memories (Squire and Bayley 2007; Dede et al. 2016; Moscovitch et al. 2016), working memory (Squire 2017; Yonelinas et al. 2024), or to material- (Lee et al. 2006; Bird and Burgess 2008; Lee et al. 2012; Mullally et al. 2012; Maguire and Mullally 2013; Kim et al. 2015; Urgolites et al. 2017), and/or process-specific roles of the hippocampal-diencephalic-cingulate network (Lee et al. 2006; Kopelman et al. 2007; Wixted and Squire 2011; Patai et al. 2015; Kafkas et al. 2017; Argyropoulos and Butler 2020). It may also clarify similarities and discrepancies in the neuropsychological profile of hippocampal vs. extra-hippocampal damage, and how the interactions of those areas support memory (Winocur 1985; Aggleton and Brown 1999; Gold and Squire 2006; Kopelman et al. 2007; Aggleton et al. 2008).

In Argyropoulos et al. (2019), we had reported a large cohort (n=38) characterised by hippocampal atrophy (which, beyond the volume reduction in the right entorhinal cortex, was focal within the medial temporal lobe) and residual amnesia due to autoimmune limbic encephalitis (aLE) – a disease studied as a “human model” of hippocampal damage (Maguire et al. 2006; Hassabis et al. 2007; Maguire et al. 2015 Aug 14; Henson et al. 2016; McCormick et al. 2016; Henson et al. 2017; McCormick et al. 2018; Lad et al. 2019; Spanò, Weber, et al. 2020) due to its focality (post-mortem and animal-model studies: Dunstan and Winer, 2006; Park et al., 2007; Khan et al., 2009; Tröscher et al., 2017) relative to ischemia/anoxia or other encephalitides (Damasio and Van Hoesen, 1985; Gitelman et al., 2001; Raschilas et al., 2002; Huang and Castillo, 2008; Heinz and Rollnik, 2015; Kelly et al., 2024). Acute clinical scans had disclosed very few abnormalities beyond the medial temporal lobes, and none in the diencephalon, unlike the significant diencephalic injury in other conditions causing hippocampal damage (Dzieciol et al. 2017; Meys et al. 2022). Post-acutely, however, we recorded whole-thalamic volume reduction and resting-state functional abnormalities, involving the cingulate. Surprisingly, it was these abnormalities, and not hippocampal volumes, that mostly explained amnesia.

Since Argyropoulos et al. (2019), methods designed to overcome the difficulties posed by conventional MRI protocols in automatically segmenting individual diencephalic nuclei have become available (Billot et al. 2020; Greve et al. 2021; Tregidgo et al. 2023; Vidal et al. 2024). In this study, we predicted that the key components of the revised Papez circuit with strong support for involvement in episodic memory (principal anterior thalamic nuclei, mammillary bodies, fornix; Aggleton et al. 2022), would show structural abnormalities, that their integrity would be associated with that of the hippocampal formation, and that these abnormalities would explain episodic memory impairment across patients. In addition to hippocampal volumes derived from gold-standard manual delineation in this cohort (Argyropoulos et al. 2019), we also estimated hippocampal/subicular subfield volumes, and whether they would show relationships with memory scores over and above diencephalic volumes. We also sought evidence consistent with a causal account, whereby the effects of hippocampal damage would be followed by those in the fornix, which would precede abnormalities in the mammillary bodies. Finally, we expected that abnormalities would be present even in anti-leucine-rich glioma inactivated (LGI1)-aLE, i.e., the type that is often the focus of many studies of hippocampal damage (Lad et al. 2019; Miller et al. 2020; Spanò, Weber, et al. 2020; Spanò, Pizzamiglio, et al. 2020), as it may present with damage and symptoms more focally and homogeneously compared with other types (van Sonderen et al. 2016; Bastiaansen et al. 2017; Finke et al. 2017; van Sonderen et al. 2017).

## Materials and Methods

### Participants

Our patient cohort (n=38; 26M:12F; age at imaging: mean=61.39; SD=13.91 years), presented in Argyropoulos et al. (2019, 2020), comprised individuals who had been diagnosed with aLE according to established criteria (Graus et al. 2016; S-Methods 1).

### MRI data acquisition

#### Structural MRI

All patients underwent structural brain imaging, along with 67 healthy controls (40M:27F; age at imaging: mean=61.23; SD=13.57 years; controls vs. patients: M:F ratio: χ^2^=0.79, p=0.374; age: t(103)=-0.06, p=0.954). T1-weighted images were acquired using a Magnetization Prepared Rapid Gradient Echo (MPRAGE) sequence (echo time=4.7ms, repetition time=2040ms, 8° flip angle, field of view=192mm, voxel size=1×1×1mm). All patients and 35/67 controls were recruited by the Memory and Amnesia Project (University of Oxford). An additional 32 structural MRI datasets of controls were made available through the Oxford Project To Investigate Memory and Aging. Although acquisition protocols did not differ (Zamboni et al. 2013), our volumetric comparisons between all controls and patients include scan source as a covariate.

#### Diffusion MRI

All diffusion data were acquired under our Memory and Amnesia Project, involving 35/35 controls (age: mean=55.14; SD=14.04 years; 25M:10F), and 37/38 patients (age: mean=61.14; SD=14.01 years; 26M:11F; controls vs. patients: M:F ratio: χ^2^=0.01, p=0.914; age: t(70)=1.81, p=0.074; age and sex were included in between-groups comparisons). We used a single-shot echo planar imaging sequence (64 slices; slice thickness=2mm, 0 gap; axially acquired; 64 directions at b=1,500s/mm^2^, repetition time=8,900ms; echo time=94.8ms; voxel size=2×2×2mm; field of view=192×192mm). One no-diffusion-weighted image at b=0s/mm^2^ was also acquired.

### Experimental Design and Statistical Analysis

#### Automated Diencephalic volumetry

We first compared controls and patients (ANCOVAs - covariates: age, sex, FreeSurfer’s estimated total intracranial volume (eTIV), scan source) on the volumes of the (left+right) mammillary bodies segmented with FreeSurfer-ScLimbic (Greve et al. 2021), and the thalamic segmentations estimated by the convolutional neural network tool developed in FreeSurfer (Tregidgo et al. 2023) (henceforth, “FreeSurfer-CNN”). FreeSurfer-ScLimbic is a deep-learning tool implemented in FreeSurfer (“mri_sclimbic_seg”; https://surfer.nmr.mgh.harvard.edu/fswiki/ScLimbic) that automatically segments several subcortical limbic structures using T1-weighted images (Greve et al. 2021) (available for 105/105 participants). FreeSurfer-CNN (https://surfer.nmr.mgh.harvard.edu/fswiki/ThalamicNucleiDTI) employs diffusion and T1-weighted MRI data (available for 37 patients and 35 controls). Using FreeSurfer 7.4.1, we first produced a bias-corrected, whole-brain structural T_1_-weighted MRI image (“norm.mgz”), a whole-brain segmentation (“aseg.mgz”) using “recon-all”, and a T_1_-based FreeSurfer segmentation (Iglesias et al. 2018), using “segmentThalamicNuclei.sh” (https://freesurfer.net/fswiki/ThalamicNuclei). We then used “TRActs Constrained by UnderLying Anatomy” (Yendiki et al. 2011; Maffei et al. 2021) to derive a fractional anisotropy volume (“dtifit_FA.nii.gz”) and a 4D volume containing the principal direction vector for each DTI voxel (“dtifit_V1.nii.gz”). The segmented nuclei comprise the central lateral, central medial, centromedian, lateral geniculate, lateral posterior, laterodorsal, limitans (suprageniculate), medial geniculate, mediodorsal lateral (parvocellular), mediodorsal medial (magnocellular), parafascicular, pulvinar anterior, pulvinar inferior, pulvinar lateral, pulvinar medial (medial segment – henceforth, “m”; lateral segment – henceforth, “l”), reuniens (medial ventral), ventral anterior, ventral anterior magnocellular, ventral lateral anterior, ventral lateral posterior, ventral posterolateral nuclei, along with an “anteroventral” segmentation, which, however, includes the anterior medial and anterior dorsal nuclei (Iglesias et al. 2018) – we will thus refer to this segmentation as the “principal anterior nuclei” instead. We applied the Holm-Bonferroni sequential correction (Holm 1979) for the number of tests (n=24) conducted (henceforth, “p-corr(24)”). We also examined whether these differences could be conceptually replicated using HIPS-THOMAS (Vidal et al. 2024) for the thalamus, and HypothalamicSubunits for hypothalamic segmentations (Billot et al. 2020; S-Methods 2). Comparing these segmentation methods is beyond the scope of this paper (Argyropoulos et al. 2025).

#### WM integrity

A standard preprocessing pipeline for single-shell single-tissue constrained spherical deconvolution was followed in MRtrix3 (Tournier et al. 2019) (S-Methods 3). Connectivity-based fixel enhancement (a threshold-free cluster-enhancement-like approach) (Smith and Nichols 2009) was involved, which used probabilistic tractography to identify structurally connected fixels that share underlying anatomy. FWE-corrected *p*-values (<0.05) were then assigned to each fixel using non-parametric permutation testing (5,000 permutations). As in recent studies (Zarkali et al. 2020; Andica et al. 2021), we selected fibre density and cross-section (FDC) as the measure of interest, since it reflects both microstructural and macrostructural changes, representing a tract’s overall ability to relay information (Raffelt et al. 2017). To conceptually replicate the findings of our fixel-based analysis, we also employed another two whole-brain automated analyses of WM integrity (voxel-based morphometry, tract-based spatial statistics), along with deterministic tractography to manually reconstruct key tracts of interest (S-Methods 4-6).

#### Structure-structure/behaviour relationships

We examined the relationship of impaired episodic memory with reduced diencephalic volumes (n=8). Volumes were residualised against the same regressors as those included in the ANCOVAs above. Correlational analyses (conducted separately for patients) employed our two composite scores - “anterograde retrieval” and “remote autobiographical memory”, which were shown to be better explained by extra-hippocampal abnormalities in our cohort (p-corr(16)). The former was the average of standardised age-scaled scores on 13 (sub)tests, on all of which the patient cohort showed impairment (Argyropoulos et al. 2019): i) visual recall: Doors and People – Shapes (Baddeley et al. 1994), Rey-Osterrieth Complex Figure Copying Test (Rey 1959) Immediate and Delayed Recall; ii) visual recognition: Doors and People – Doors (Baddeley et al. 1994), Warrington Topographical Memory test (Warrington 1996); iii) verbal recall: Wechsler Memory Scale III – Logical Memory I/II, Word List I/II (Wechsler 1997), Doors and People – People (Baddeley et al. 1994); iv) verbal recognition: Doors and People – Names (Baddeley et al. 1994), Wechsler Memory Scale III – Word List II Recognition (Wechsler 1997), Warrington Recognition Memory Tests – words (Warrington 1984). The latter was the sum of autobiographical memory scores for Childhood and Early Adulthood (Kopelman et al. 1989; see Argyropoulos et al. (2019) for the reasons why the ’Recent’ phase scores were disregarded). These correlations were then iterated, after decomposing the anterograde composite score into verbal/visual recall/recognition (p-corr(40)) and also by examining laterality-specific relationships (p-corr(80)).

We further estimated hippocampal/subicular subfield volumes, in order to ascertain whether these, unlike our gold-standard manually delineated hippocampal volumes (Argyropoulos et al. 2019), strongly predicted episodic memory scores (a fortiori, over and above the diencephalic volumes here). We did so, while deliberately disregarding correction for multiple correlational tests, as well as the fact that 1mm^3^ -resolution MRIs may be too low-quality for automated subfield volume estimates (Wisse et al. 2021). We first used FreeSurfer’s “Subregions Segmentation” (Iglesias et al. 2015) to derive subfield segmentations of the hippocampal formation. This tool (“segment_subregions”; https://surfer.nmr.mgh.harvard.edu/fswiki/SubregionSegmentation) was used for segmentation of MR images pre-processed through the FreeSurfer “recon-all” pipeline. It outputs segmentations for the right/left hippocampal formation: amygdala transition area, CA1/CA3/CA4 head/body, subiculum head/body, presubiculum head/body, molecular layer head/body, granule cell and molecular layer of the dentate gyrus head/body, hippocampal fissure, parasubiculum, tail, and fimbria. We excluded the fimbria (WM) from our volumetric comparisons. A series of ANCOVAs (covariates: age, sex, scan source, and eTIV) on raw total (left+right) volumes showed that all subfields, apart from the parasubiculum and the hippocampal fissure, were atrophic (15%-26%; p-corr(18)<0.05; *η^2^_p_*=0.11-0.36). We then used the reduced (left+right) subfield volumes (residualised against age, sex, scan source, and eTIV) that showed the (numerically) highest correlation coefficient with a composite memory score of interest as a control variable in partial correlation analyses of the relationships between diencephalic volumes and memory scores. We also conducted a series of mediation analyses, to investigate whether any relationships observed between these hippocampal/subicular subfield volumes (predictor) and episodic memory scores (outcome) were mediated by the affected diencephalic volumes across patients. Mediation analyses rely on the assumption of causality between the predictor and the mediator (Judd and Kenny 1981; Baron and Kenny 1986; MacKinnon et al. 2007). A default number of 5,000 bootstrap samples was used to calculate the standard errors and confidence intervals.

For the purposes of examining structure-structure/behaviour relationships, we used a tract-of-interest analysis. Major tracts (n=70/72 totally generated, excluding the inferior cerebellar peduncle, as inferior cerebellar slices were clipped from the field of view) were automatically delineated on the WM fibre orientation distribution template using TractSeg (Wasserthal et al. 2018). Delineated fibre tracts were then converted into fixel masks after using the “tck2fixel” command in MRtrix3 (Tournier et al. 2019), and subsequently thresholding this fixel image to create a binary tract fixel mask. We extracted mean tract FDCs, and, for the purposes of reducing the number of tracts and hence the number of tests conducted, we collapsed across hemispheres, and also derived a single mean FDC for the corpus callosum from the 7 available (“CC 1-7”). We then compared controls and patients in a series of ANCOVAs (covariates: age, sex; p-corr(33)) to identify the tracts affected in aLE. We then entered the mean FDC (residualised for age and sex) of the affected tracts (n=25) into correlational analyses with the volumes (residualised for age, sex, scan source, and eTIV) of the mammillary body and the hippocampal formation (p-corr(50)), to identify tracts whose integrity was associated with that of the mammillary body and the hippocampal formation. We predicted that the integrity of the fornix would show strong relationships with that of both the hippocampus and the mammillary bodies across patients, and that the effects of hippocampal volume (predictor) on mammillary body volume (outcome) would be mediated by the FDC of the fornix, given our hypothesis that any reduction in the volume of mammillary bodies would follow as a knock-on effect of damage in the hippocampal formation and subsequently in the fornix. A default number of 5,000 bootstrap samples was used to calculate the standard errors and confidence intervals.

We ran a series of correlations between the mean FDC (residualised) of the affected tracts (n=25) and anterograde retrieval / remote autobiographical memory (p-corr(50)). In a post-hoc, exploratory fashion, we examined whether the integrity of the fornix also predicted Recollection and/or Familiarity for Faces, Scenes, and/or Words across the 8 aLE patients who had completed custom-made recognition memory tasks, reported in Argyropoulos et al. (2022). Mean FDC (residualised), was entered into a linear mixed effects model (using “lmer” in R), with fully factorial fixed effects of mean FDC, Process, Material-Type, Paradigm, and Hemisphere. A single random intercept across participants was used. The results were reported in terms of Type III ANOVA using Satterthwaite’s method for adjusting degrees of freedom. We then iterated this analysis, separately for Process (Recollection, Familiarity: p-corr(2)), and Process-and-Material-Type (Faces/Scenes/Words x Recollection/Familiarity: p-corr(6)).

Finally, comparisons with healthy controls on diencephalic volumes and mean FDC per tract were iterated for the subgroup of n=14 LGI1-aLE patients (Argyropoulos et al. 2019) in our cohort. Comparisons with other autoantibody-related subsets of our cohort were not meaningful or feasible- e.g., at least some of the 10/38 seronegative patients and some of the 7/38 that were found positive for antibodies targeting the voltage-gated potassium channel complex may have had LGI-1 aLE.

## Results

Our cohort showed volumetric reduction in the mammillary bodies (7%; *η^2^_p_*=0.09), the principal anterior nuclei (36%; *η^2^_p_* =0.33), the medial (m) (31%; *η^2^_p_*=0.32) and lateral pulvinar (13%; *η^2^_p_*=0.15), the medial mediodorsal (22%; *η^2^_p_*=0.23) and lateral mediodorsal nuclei (22%; *η^2^_p_*=0.19), but also the laterodorsal (21%; *η^2^_p_*=0.17) and limitans-suprageniculate segmentation (20%; *η^2^_p_*=0.14; all ANCOVAs: p-corr(24)<0.05; Fig. 1;Table S1). These differences were conceptually replicated with HIPS-THOMAS and HypothalamicSubunits (Table S2).

**Fig. 1:**
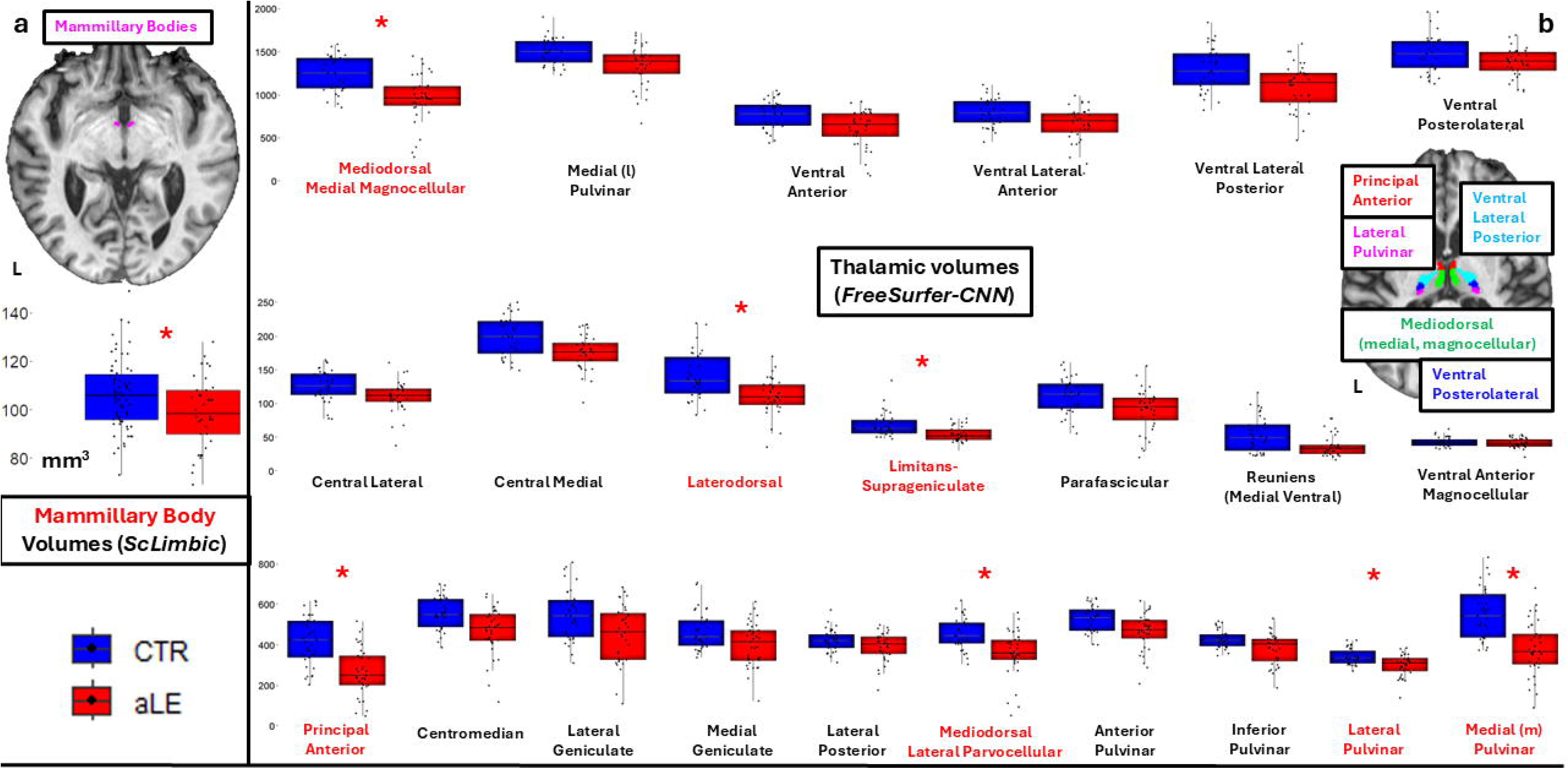
volumes for (**a)** mammillary bodies and (**b)** thalamic nuclei; **key:** lines in boxplots=medians; bottom of box=25th %ile; top of box□=□75th %ile; upper,lower whiskers=scores outside the middle 50; whiskers□=□1.5×interquartile range; *: p-corr(Holm 1979)<0.05; aLE: autoimmune limbic encephalitis patients; CTR: healthy controls; L/R: Left/Right (Hemisphere).

Of the affected (n=8) nuclei, the laterodorsal, medial (m) pulvinar, and mammillary bodies correlated with the volume of the manually delineated hippocampal formation (r=0.47-0.52, p-corr(8)=0.007-0.016; Fig. 2f-h). Of these 8 diencephalic nuclei, the mammillary bodies and the laterodorsal nuclei showed a volumetric relationship with anterograde retrieval (Fig. 2a-b), while remote autobiographical memory was associated with the volumes of the laterodorsal, limitans-suprageniculate, and medial (m) pulvinar segmentations (Fig. 2c; r=0.46-0.62, p-corr(16)<0.05; rest: r=0.19-0.43, p-corr(16)≥0.137). None of these volumes was associated with the semantic aspect of remote memories (r≤0.18; p≥0.339). We then iterated these analyses, after decomposing the anterograde composite score into verbal/visual recall/recognition scores, also examining hemisphere-specific volumes.

**Fig. 2:**
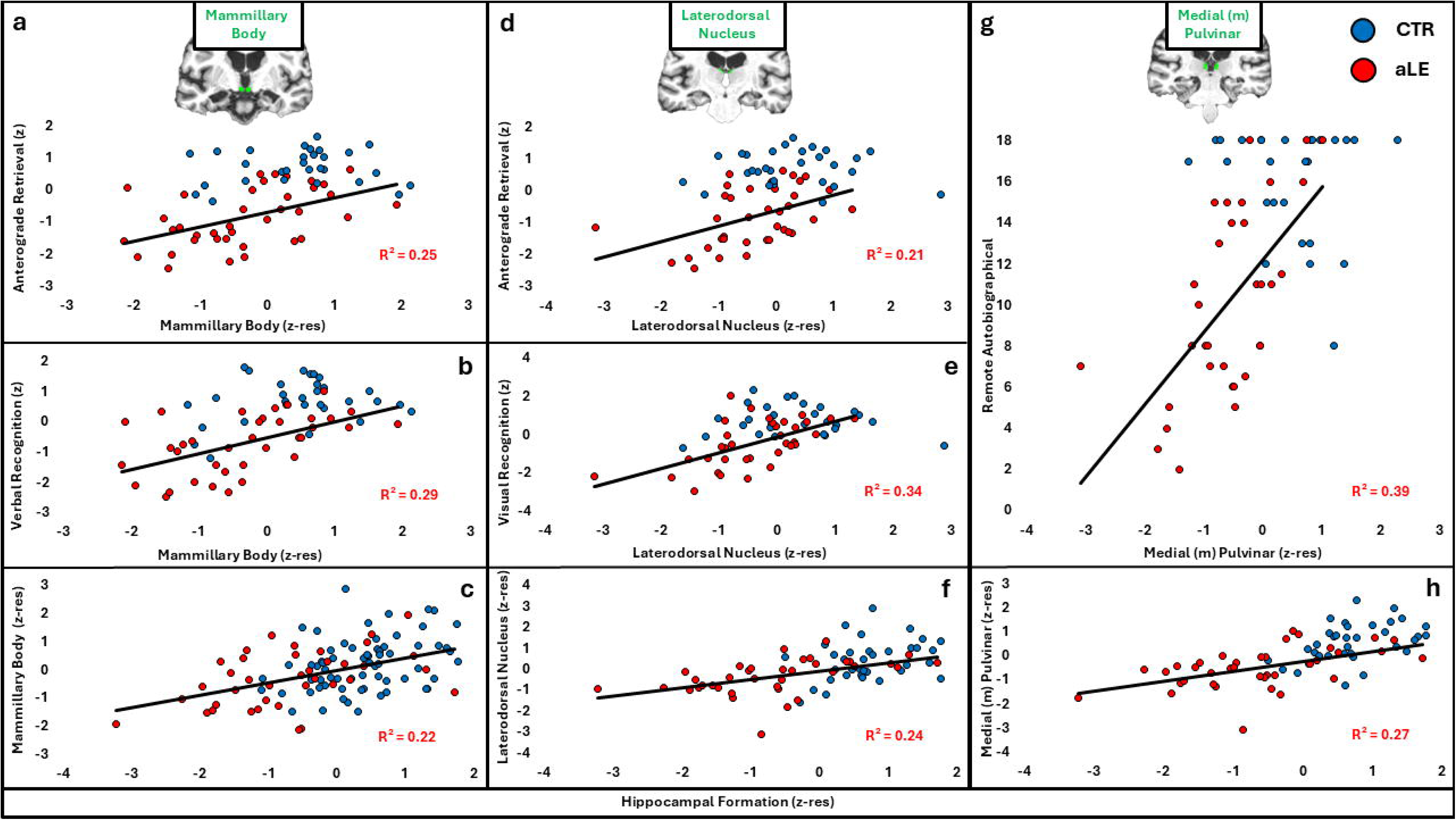
a-c: relationships of mammillary body volumes with (a) anterograde retrieval composite score, (b) verbal recognition composite score, and (c) hippocampal formation volume; d-f: relationships of laterodorsonal nucleus volumes with (d) anterograde retrieval composite score, (e) visual recognition, and (f) hippocampal formation volume; g-h: relationships of medial (m) pulvinar volume with (g) remote autobiographical memory and (h) hippocampal formation volume. Data from healthy controls are not used in the calculation of the R^2^; **key:** aLE: autoimmune limbic encephalitis patients; CTR: healthy controls; z-res: standardised residuals.

**Verbal recognition memory** correlated with mammillary body volumes (r=0.54, p-corr(40)=0.021; Fig. 2d; rest: r≤0.39; p-corr(40)≥0.485), driven by the left mammillary body (r=0.61, p-corr(80)=0.006; rest: r≤0.43; p-corr(80)≥0.497). While this volume correlated, at uncorrected levels, with verbal/visual recall and visual recognition scores (r=0.41-0.53, p<0.05, p-corr(80)≥0.054), partial correlation analyses showed the relationship with verbal recognition held over and above visual recall/recognition (r=0.41-0.45, p=0.008-0.016), and, marginally, verbal recall (r=0.30, p=0.075). **Visual recognition memory** correlated with the volume of the laterodorsal nucleus (r=0.58, p-corr(40)=0.009; Fig. 2e), and marginally with medial (m) and lateral pulvinar volumes (r=0.49-0.51, p-corr(40)=0.064-0.086; rest: r≤0.42, p-corr(40)≥0.346). The relationship with the laterodorsal nucleus held over and above those with the medial (m) and lateral pulvinar (r=0.35-0.42, p=0.015-0.040). This was driven by the left laterodorsal nucleus (r=0.60, p-corr(80)=0.012; rest: r≤0.52, p-corr(80)≥0.107). While this volume also correlated with visual/verbal recall and verbal recognition (r=0.43-0.51, p=0.001-0.008), its links with visual recognition held over and above the rest (r=0.40-0.46, p=0.009-0.019). The relationship of the medial (m) pulvinar volume with **remote autobiographical memory** was driven by the left medial (m) pulvinar (r=0.65, p-corr(80)=0.004; left limitans-suprageniculate: r=0.58, p-corr(80)=0.031; right medial (m) pulvinar: r=0.57, p-corr(80)=0.046; left laterodorsal nucleus: r=0.55, p-corr(80)=0.075; rest: r≤0.52, p-corr(80)≥0.131). Its link with remote autobiographical memory held over and above these volumes (r≥0.38; p≤0.033). Importantly, these diencephalic relationships with memory held over and above (partial correlation) the total volume of the manually delineated hippocampal formation, as well as the strongest observable correlations of hippocampal/subicular subfield volumes with memory scores; in fact, diencephalic integrity fully mediated the effects of hippocampal volumes on memory scores (Table S3). No relationships between diencephalic volumes and **verbal/visual recall** survived correction(|r|≤0.53; p-corr(80)≥0.054).

A whole-brain fixel-based analysis showed reduced FDC in the fornix, cingulum, mammillothalamic tract, and internal capsule. However, abnormalities extended to the anterior commissure, the corpus callosum and its radiations, and the superior cerebellar peduncle (Fig. 3a). These were broadly replicated with voxel-based morphometry, tract-based spatial statistics, and tract reconstruction using deterministic tractography (Fig. S1-3).

**Fig. 3:**
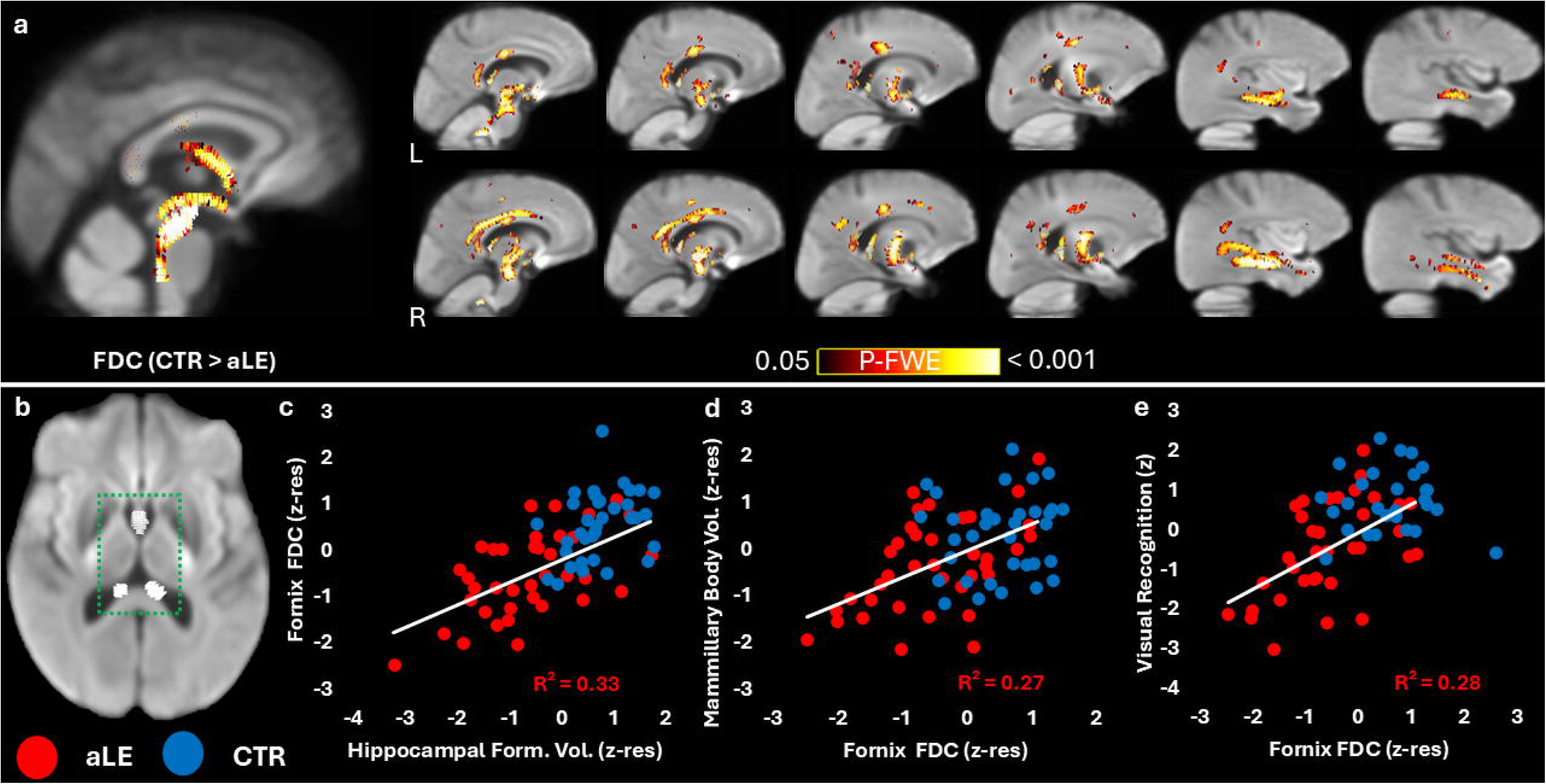
a: Fixel-based analysis comparisons between controls and patients on FDC. Clusters survive whole-brain threshold-free cluster-enhancement-correction (p-FWE<0.05); b: binary fornix fixel mask; **c-e:** relationship of mean fornical FDC across patients with (c) hippocampal formation volumes, (d) mammillary body volumes, and (e) visual recognition memory composite scores (Argyropoulos et al. 2019) (healthy control data are not used in the calculation of the R^2^); **key:** aLE: autoimmune Limbic Encephalitis; CTR: healthy controls; FDC: Fiber Density and Cross-section; L/R: left/right hemisphere; z-res: standardised residuals.

Of the tracts compared between groups and found to show reduced FDC in patients (n=25; Table S4), the fornix showed the largest effect size (*η^2^_p_*=0.32), followed by the superior cerebellar peduncle (*η^2^_p_*=0.32; rest: *η^2^_p_*=0.11-0.21). The fornix was also the only tract with reduced FDC to show a relationship, across patients, with hippocampal (r=0.57, p-corr(50)=0.011) and mammillary body volumes (r=0.52, p-corr(50)=0.048; rest: |r|<0.48, p-corr(50)>0.132). Corroborating the idea that these abnormalities occur as knock-on effects following hippocampal damage (Argyropoulos et al. 2019), the relationship between the hippocampal formation and the mammillary bodies was at least partly mediated by the integrity of the fornix (Fig.S4).

No relationship between mean tract FDC and the two composite scores (anterograde retrieval, remote autobiographical memory) survived correction for multiple tests (|r|<0.53, p-corr(50)<0.080), and the integrity of the fornix correlated with these scores only at uncorrected levels (remote autobiographical memory: r =0.36, p=0.039; anterograde retrieval: r=0.35, p-unc=0.034). The relationship with the latter was driven by visual recognition (r=0.53, p=0.001; verbal recognition: r=0.30, p=0.071; verbal recall: r=0.34, p=0.042), which held over and above verbal recall (r=0.43, p=0.012), verbal recognition (r=0.47, p=0.005), visual recall (r=0.51,p=0.002), and remote autobiographical memory (r=0.40,p=0.027). This pattern was corroborated by a series of exploratory analyses on the behavioural data from 8/38 aLE patients who had participated in custom-made recognition memory tasks dissociating recollection from familiarity (Argyropoulos et al. 2022). Fornical FDC was associated with familiarity for topographical scenes, not with face or word familiarity, or recollection (Table S5).

Consistent with the whole cohort, the subgroup (n=14/38) of LGI1-aLE patients showed atrophy in the anterior, mediodorsal, and pulvinar nuclei, and compromised WM integrity in the fornix and beyond (Tables S6-8).

## Discussion

In a large patient cohort (n=38) characterised by hippocampal atrophy and residual episodic memory impairment, we showed compromised integrity in specific diencephalic nuclei and WM tracts. These abnormalities explained aspects of amnesia over and above hippocampal volumes.

While whole-thalamic structural abnormalities have been demonstrated in aLE, and even specifically in LGI1-aLE (Argyropoulos et al. 2019; Qiao et al. 2025), there is hardly any evidence for nucleus-specific abnormalities in this condition. Here, the extent of the anterior thalamic atrophy (21%-36%) was comparable to, if not larger than that of the manually delineated hippocampal formation (22%-25%) (Argyropoulos et al. 2019), or the hippocampal-subicular subfields (15%-26%). Despite the evidence for atrophy in the anterior thalamus in medial temporal lobe epilepsy (Mueller et al. 2010; Tung et al. 2022) and its role in memory (de Bourbon-Teles et al. 2014; Sweeney-Reed et al. 2021; Aggleton et al. 2022), no relationships survived multiple-testing correction. This may reflect false negatives, lack of granularity in the segmentation of the anterior principal nuclei, or stronger involvement of the anteromedial and anterodorsal nuclei.

Mediodorsal atrophy (10%-22%) is also seen in temporal lobe epilepsy (Turski et al. 1986; Shimosaka et al. 1992). Evidence for projections from the perirhinal cortex and the adjacent amygdala (Graff-Radford et al. 1990; Parkin et al. 1994), task-based fMRI studies (Kafkas and Montaldi 2014), and neuropsychological research (Carlesimo et al. 2011) advocate for dissociations between mediodorsal and anterior nuclei in familiarity vs. recollection, respectively. However, we found no relationships with memory scores, and there is growing recognition that these nuclei primarily support executive aspects of memory (Mair et al. 2015; Perry et al. 2021; Wolff and Halassa 2024).

Atrophy in the laterodorsal nucleus (∼21%) dovetails with its similarities in neocortical connectivity with the anterior thalamic nuclei (Robertson and Kaitz 1981; Jones 2012) and its inclusion in the anterior nuclear complex (Morel et al. 1997; Krauth et al. 2010). Consistent with its volumetric relationship with the hippocampal formation here, it has direct interconnections with the hippocampal formation (Aggleton et al. 1986; Van Groen and Wyss 1992; Vertes et al. 2007; Varela et al. 2014). Like the anterior thalamic nuclei, it is reciprocally connected with the posterior cingulate (Horikawa et al. 1988; Aggleton et al. 2014), wherein our patients showed reduced resting-state activity (Argyropoulos et al. 2019). Its volume predicted visual recognition memory, over and above any hippocampal-subicular subfield volumes. This dovetails with its proposed involvement in visuospatial memory and the head direction signals it provides to the hippocampal formation (Warburton et al. 1997; van Groen et al. 2002; Perry and Mitchell 2019). Less is known about it in humans (Saunders et al. 2005; Cipolotti et al. 2008), or its role within the extended hippocampal system (Aggleton and Brown 1999; Aggleton et al. 2010; Aggleton et al. 2022). Its relationship with visual recognition memory over and above visual recall appears consistent with evidence that it contributes to familiarity (Cipolotti et al. 2008). However, our recognition tests involved photographs of doors and topographical scenes, whereas recall involved arbitrary shapes.

We also saw medial (m) pulvinar atrophy (∼31%). Whereas the rest of the pulvinar mainly connects with parietal and occipital cortices (Gutierrez et al. 2000), the medial pulvinar is reciprocally connected with the temporal lobe (Burton and Jones 1976; Mauguière and Baleydier 1978; Yeterian and Pandya 1985; Insausti et al. 1987; Baleydier and Morel 1992; Webster et al. 1993). Apart from perirhinal and entorhinal projections to the medial pulvinar via the temporopulvinar bundle (Saunders et al. 2005), the non-fornical projection from the subicular cortex to the laterodorsal nucleus passes through the medial pulvinar, with some additional termination there (Aggleton et al. 1986). In humans, the medial pulvinar is strongly connected (structurally, functionally) with the hippocampus (Guye et al. 2006; Rosenberg et al. 2006; Rosenberg et al. 2009; Catenoix et al. 2011; Morgan et al. 2015; Voets et al. 2015; Zheng et al. 2018; Segobin et al. 2019; Capecchi et al. 2020; Soulier et al. 2023) and the posterior cingulate (Baleydier and Mauguiere 1985; Shibata and Yukie 2003; Parvizi et al. 2006), wherein our patients showed reduced resting-state activity. It has also shown strong involvement in genetic frontotemporal dementia (Soskic et al. 2025), and, consistent with the strong link we observed with remote autobiographical memory, it is embedded within the default-mode network (Li et al. 2021), with its integrity being crucial for autobiographical retrieval (Philippi et al. 2015).

Despite our study’s cross-sectional nature, the evidence above and our mediation analyses support the possibility that these abnormalities reflect knock-on effects, akin to those in animal models (Machado et al. 2008; Meng et al. 2014; Meng et al. 2016). Of the few longitudinal MRI studies on aLE, Wagner et al. (2015) reported volume reduction from the acute to the chronic stage in the medial temporal lobe, the basal ganglia, but also the thalamus, albeit with no further details.

Given its connectivity with the hippocampal formation and the prefrontal cortex (Yanagihara et al. 1985; Vertes 2006; Vertes et al. 2007; Varela et al. 2014), the absence of atrophy in the medioventral–reuniens segmentation is hard to interpret, as is the reduction in the limitans-suprageniculate. Despite the evidence for the connectivity of the latter with the prefrontal (Preuss and Goldman-Rakic 1987; Burman et al. 2011) and perirhinal cortices (Kimura et al. 2003; Tomás Pereira et al. 2016), there is little information on abnormalities in hippocampal damage (Lee et al. 2020) or on its involvement in memory (Kim et al. 2020). Given the very small volumes of these two segmentations, and that they have only been generated with FreeSurfer-CNN, replication is required in other cohorts.

All WM analyses disclosed abnormalities in the fornix, complementing the atrophy in the mammillary bodies and the principal anterior nuclei - projections to the latter almost exclusively involve the fornix (Saunders et al. 2005). Of the affected tracts, mammillary body volumes exclusively correlated with the fornix, consistent with recent work in unilateral mesial temporal sclerosis (Mojica et al. 2026). A mediation analysis supported the plausibility of a causal link, whereby hippocampal damage compromises the fornix, which in turn compromises the mammillary bodies. There is evidence consistent with Wallerian degeneration of the fornix following hippocampal damage (Baldwin et al. 1994; Fernandez et al. 2022) and of the mammillary bodies following fornical transections (Zola-Morgan et al. 1989; Loftus et al. 2000), while fornix damage does not produce retrograde degeneration in its fibres from the temporal lobe (Daitz and Powell 1954; Saunders and Aggleton 2007). Mammillary body volumes correlated strongly with verbal recognition, over and above hippocampal volumes and, marginally, verbal recall. Correlations of fornical integrity (FDC) with memory scores were only observed at uncorrected levels, and only with visual recognition memory (Argyropoulos et al. 2019) and familiarity estimates for topographical scenes (Argyropoulos et al. 2022). This is not consistent with studies in healthy participants (Nestor et al. 2007; Rudebeck et al. 2009; Hartopp et al. 2019; Metzler-Baddeley et al. 2019; Coad et al. 2020) or patients (Carlesimo et al. 2007; Tsivilis et al. 2008; Vann et al. 2009) showing selective involvement of the fornix/mammillary bodies in recall/recollection (Aggleton and Brown, 1999; Montaldi and Mayes, 2010; Aggleton et al., 2023 – however, see: Calabrese et al., 1995; D’Esposito et al., 1995; Park et al., 2000; Gold and Squire, 2006; Poreh et al., 2006; Cipolotti et al., 2008). Regardless, our findings cannot reflect a replication failure, given substantial differences in our patients’ aetiological and neuropathological profiles.

Abnormalities in the cingulum (captured with all methods) are consistent with the atrophy in the anterior nuclei, since the cingulum involves projections of these nuclei to the hippocampal formation (Aggleton et al. 2022). They also dovetail with the reduced resting-state activity and hippocampal connectivity seen in the cingulate and retrosplenial cortices (Argyropoulos et al. 2019), since the cingulum comprises reciprocal connections between the anterior thalamic nuclei and the cingulate and retrosplenial cortices (Domesick 1970; Shibata 1993a; Shibata 1993b; Heilbronner and Haber 2014; Bubb et al. 2018; Bubb et al. 2020).

Abnormalities were also seen in the mammillothalamic tract and the internal capsule. The latter carries projections from the hippocampal formation and entorhinal cortex to the principal anterior and laterodorsal nuclei (Aggleton et al. 1986; Saunders et al. 2005; Dillingham et al. 2015), both of which were atrophied in our patients.

In agreement with recent evidence for whole-brain structural connectivity and aberrant resting-state functional connectivity across several large-scale networks in LGI1-aLE (Heine et al. 2018; Qiao et al. 2020; Krohn et al. 2025), we saw abnormalities beyond the hippocampal-diencephalic-cingulate network, especially in the corpus callosum and its radiations, the superior cerebellar peduncle, and the external capsule. The last two dovetail with the atrophic mediodorsal thalamus: the lateral cerebellar nucleus reaches the lateral mediodorsal nucleus via the superior cerebellar peduncle (Çavdar et al. 2014), and the external capsule carries entorhinal-perirhinal projections to the mediodorsal nuclei (Saunders et al. 2005). Callosal abnormalities have been reported in non-human primates following hippocampal damage (Meng et al. 2014; Payne et al. 2017), and in patients with hippocampal sclerosis (Raffelt et al. 2017). Although the hippocampus does not send inter-hemispheric projections via the corpus callosum, the cortical regions that are interconnected with the hippocampus do (Meng et al. 2014; Meng et al. 2018).

Overall, our work urges caution in assuming “focal hippocampal damage”, even in patients with conditions studied as “human models” of such damage. It also underscores the possibility that extra-hippocampal abnormalities that may occur as knock-on effects of hippocampal damage, such as those in specific diencephalic nuclei (even beyond the principal anterior nuclei or the mammillary bodies), may strongly predict amnesia, over and above hippocampal integrity.

## Acknowledgments

We are very grateful to the participants who took part in this study.

- We also thank Drs Clare Loane, Adriana Roca-Fernandez, Carmen Lage-Martinez, and Sarosh R Irani, for participant recruitment and data acquisition (Argyropoulos et al. 2019).

- We are grateful to Drs Henry Tregidgo, Manojkumar Saranathan, and Arkiev D’Souza, for their responses to our questions on the use of FreeSurfer-CNN, HIPS-THOMAS, and MRtrix3, respectively.

- We are also grateful to the anonymous reviewers of earlier versions of this manuscript for their helpful recommendations.

