## Supplemental Methods, Figures, and Tables for "Diencephalic integrity explains aspects of hippocampal amnesia"

### Supplementary Methods

#### S-Methods 1: Patient Cohort details

The most common autoantibody (n=14/38) was anti-leucine-rich glioma inactivated (anti-LGI-1); 10/38 were seronegative, but may have had LGI-1 aLE, despite that anti-LGI-1 was not detectable in routine clinical practice at the time of screening (Graus et al. 2018); 2/38 were positive for anti-glutamic acid decarboxylase autoantibody; 1/38 presented with autoantibodies characteristic of paraneoplastic encephalitis; 4/38 presented with dual seropositive antibodies of contactin-associated protein-like 2 and LGI-1, and 7/38 were found positive for antibodies targeting the voltage-gated potassium channel complex, with no further information on the specific antibodies. After being assessed by CRB (neurologist), they were recruited for neuropsychological assessment and research scans in the post-acute stage of the disease (median=5.41; IQR=5.36 years since symptom onset). They were all fluent in English (37 native speakers; 1 non-native speaker) and had undergone MRI at the time of initial clinical presentation as well as neuropsychological assessment at the Russell Cairns Unit, Oxford, UK (2013–8). Most of them had been treated acutely with immunotherapy (n=31/38), and had shown hippocampal abnormalities (in signal, volume, and/or diffusion) on clinical MRI conducted acutely (n=34/38). Beyond the hippocampus, 6/38 showed abnormalities in the amygdala, 1/38 in the entorhinal cortex, 1/38 in the parahippocampal cortex, 1/38 in the caudate, and 4/38 showed mild microangiopathic changes (commonly found with aging). None of them had a history of neurologic or psychiatric disorders that could have resulted in cognitive impairment.

#### S-Methods 2: Conceptual replication of automated diencephalic segmentation

HIPS-THOMAS (<https://github.com/thalamicseg/thomas_new>) is a variant of the THalamus Optimized Multi Atlas Segmentation method (Su et al. 2019). It involves a HIstogram-based Polynomial Synthesis as a preprocessing step, which outputs WM-nulled-like image contrast from standard T_1_-weighted structural MRI, by means of polynomial approximation. It is motivated by the observation that WM-nulled-MPRAGE images provide higher intra-thalamic contrast(Su et al. 2019) and generate more accurate segmentation (Umapathy et al. 2022) relative to T_1_-weighted MRIs. HIPS-THOMAS has been shown to surpass the T_1_-based FreeSurfer segmentations(Iglesias et al. 2018) in accuracy (Williams et al. 2024). The segmented thalamic nuclei (HIPS-THOMAS v.1) comprised the anteroventral, ventral anterior, ventral lateral anterior, ventral lateral posterior, ventral posterolateral, pulvinar, lateral geniculate, medial geniculate, centromedian, mediodorsal-parafascicular, and habenular nuclei. We used the “-big” option, which is recommended for cases with enlarged ventricles, given the age of our participants. The HypothalamicSubunits tool (Billot et al. 2020) (<https://surfer.nmr.mgh.harvard.edu/fswiki/HypothalamicSubunits>) segments the hypothalamus and its associated subunits in 3D T1-weighted scans, producing segmentation maps for the left/right inferior and superior tubular, anterior-inferior, anterior superior, and posterior hypothalamus (wherein the mammillary bodies reside).

#### S-Methods 3: Fixed-based analysis: preprocessing

Fixel-based analysis has been developed to address challenges faced by tract-based spatial statistics and other voxel-based analyses of WM integrity, such as the lack of specificity of individual fibre populations within each voxel, or the fact that traditional measures (e.g., fractional anisotropy) are difficult to interpret in regions involving crossing fibres (Raffelt et al. 2015; Raffelt et al. 2017). Unlike voxels, “fixels” reflect an individual **fi**bre population within a voxel, and directly relate to the underlying WM anatomy. They are typically derived from WM fibre orientation distributions, which are computed by constrained spherical deconvolution  techniques (Tournier et al. 2007). Fixel-based analysis thus enables the identification of specific fiber pathways, even within regions that contain crossing fibers. By enabling the estimation of the number of axons and the axon diameter within a voxel, the inferences drawn from these analyses may be more biologically meaningful (Raffelt et al. 2017).

Preprocessing (<https://mrtrix.readthedocs.io/en/dev/fixel_based_analysis/st_fibre_density_cross-section.html>) involved: denoising and Gibbs ringing removal; correction for motion and eddy current-induced distortions; bias field correction to eliminate low frequency intensity inhomogeneities across the image; global intensity normalisation (Raffelt et al. 2017) in order for the absolute amplitudes of the fibre orientation distributions to be comparable between participants; computation of single-shell response functions representing single-fibre WM from the data, followed by a single unique (average) response function to perform spherical deconvolution of all subjects (Raffelt et al. 2012); upsampling to a recommended voxel size of 1.25 × 1.25 × 1.25 mm in order to increase anatomical contrast and improve downstream template building, registration, tractography and statistics; computation of upsampled brain mask images; Fibre Orientation Distribution estimation, using constrained spherical deconvolution with the unique WM response function calculated above; generation of a study-specific (based on 30 subjects) unbiased template of Fibre Orientation Distribution, by averaging the fibre orientation distributions from 30/72 subjects (15 controls and the 15 patients that showed the smallest volume reduction in the thalamus and the hippocampal formation, in order to avoid patients with excessive abnormalities compared to the rest of the population, in keeping with the MRtrix3 instructions above); nonlinear registration of all subjects’ fibre orientation distribution images to the template (Raffelt et al. 2011); warping all subject masks into template space and computing a template mask as the intersection of all subject masks in template space, to ensure analysis is confined to voxels that contain data from all subjects; segmentation of fixels from the template, resulting in a fixel mask defining the fixels on which statistical analysis will be conducted; warping fibre orientation distribution images to template space; segmentation of each lobe of fibre orientation distributions to estimate the number and orientation of fixels in each voxel and their apparent fibre density; re-orienting the fixels of all subjects in template space, using the local transformation at each voxel, determined by the warps above; assigning subject fixels to template fixels; computing the log of the fibre cross-section metric, to ensure that data are centred around zero and normally distributed; computation of a combined metric of fibre density and cross-section (Raffelt et al. 2017); whole-brain fibre tractography on the fibre orientation distribution template, which involves statistical analysis using connectivity-based fixel enhancement, which exploits local connectivity information derived from probabilistic fibre tractography (20 million streamlines), which acts as a neighbourhood definition for a threshold-free cluster-enhancement; applying “Spherical-deconvolution-Informed Filtering of Tractograms” (Smith et al. 2013) to reduce biases in the tractogram densities and the number of streamlines to 2 million; generation of a fixel-to-fixel connectivity matrix, based on the whole-brain streamlines tractogram; smoothing of fixel data based on the sparse fixel-to-fixel connectivity matrix.

#### S-Methods 4: Conceptual replication of fixel-based analysis with Voxel-based Morphometry

Originally developed and commonly used to identify local changes in grey matter in T_1_-weighted structural MRI (Ashburner and Friston 2000; Good et al. 2001), voxel-based morphometry can also be used to examine WM integrity (Bray et al. 2015; Pezzoli et al. 2018). An obvious limitation is its reduced sensitivity, given that WM structures involve large homogeneous regions with limited intensity changes (Kurth et al. 2015), as well as that WM T_1_ signal intensity may be disclosed as normal, even in the presence of severe damage (Filippi et al. 2001; Büchel et al. 2004; Padovani et al. 2006). It has been noted, however, that, relative to diffusion MRI analyses, WM- voxel-based morphometry may have the advantage of more limited susceptibility to artifacts caused by motion or crossing fibers (Le Bihan et al. 2006; Pezzoli et al. 2018). We thus used WM- voxel-based morphometry as a first crude step in identifying WM damage in our patients. Using the Statistical Parametric Mapping software (SPM12 v7771; <http://www.fil.ion.ucl.ac.uk/spm/software/spm12>) in Matlab R2023a, T_1_-weighted structural MRIs were reoriented to have the anterior commissure as the same point of origin, and were bias-corrected to remove intensity non-uniformities. The unified segmentation procedure (Ashburner and Friston 2005), was used to segment images into grey matter, WM, and cerebrospinal fluid. We then employed the “diffeomorphic anatomical registration through the exponentiated lie algebra” (DARTEL) toolbox for all participants’ grey matter, WM, and cerebrospinal fluid to register images across subjects and generate study-specific WM templates (Ashburner 2007). The WM templates were affine-registered to the tissue probability maps in MNI (Montreal Neurological Institute, Quebec, Canada) space (voxel size: 1 mm^3^ isotropic). Voxel values in the tissue maps were modulated by the Jacobian determinant (calculated during spatial normalization). Modulated images, reflecting WM volume, were smoothed using a standard Gaussian filter (4 mm full-width at half maximum), to retain spatial specificity. We compared WM volume between groups (CTR>aLE), including age, sex, scan source, and total intracranial volume. We used Threshold-Free Cluster Enhancement (TFCE)(Smith and Nichols 2009) to correct for multiple comparisons (p-FWE<0.05; based on 5,000 permutations), given that this correction method was the only one commonly available across all three methods used (voxel-based morphmoetry, tract-based spatial statistics, fixel-based analysis). For the voxel-based morphometry analysis, we used the TFCE tool developed for SPM by Dr Christian Gaser (<https://github.com/ChristianGaser/tfce>).

#### S-Methods 5: Conceptual replication of fixel-based analysis with Tract-Based Spatial Statistics (TBSS)

TBSS is a widely used method of identifying voxel-wise changes in WM integrity in diffusion MRI data, overcoming limitations of WM- voxel-based morphometry of T_1_-weighted structural MRI and voxel-based morphometry-style analyses of diffusion MRI data (Smith et al. 2006). We used TBSS to examine differences between CTRs and aLE patients in Fractional Anisotropy, Axial Diffusivity, and Radial Diffusivity. FA refers to the fraction of water diffusion that is anisotropic, i.e., directionally dependent. AD (“parallel diffusivity”), pertains to the water diffusion rate along the primary, longitudinal axis of diffusion, i.e., along, or parallel to the axon. RD (“perpendicular diffusion”) refers to the magnitude of water diffusion perpendicular to fiber tracts. A standard preprocessing pipeline was used (<https://fsl.fmrib.ox.ac.uk/fsl/docs/#/diffusion/tbss>). FA images were created from the diffusion data, using FMRIB's Diffusion Toolbox (<https://fsl.fmrib.ox.ac.uk/fsl/docs/#/diffusion/index>). Non-linear registration of all images was applied into standard space. The data were corrected for the effects of head movement and eddy currents (“eddy_correct”). A brain mask was created (“bet”) on the b=0 (no diffusion weighting) image and a diffusion tensor model was fitted (“dtifit”). FA images were eroded slightly and the end slices were zeroed, to remove likely outliers from the diffusion tensor fitting. Nonlinear registration was then applied, aligning all FA images to a 1 mm^3^ isotropic standard space, using the “FMRIB58_FA” as the target. This involved carrying out one registration per subject (“-T” flag). All subjects’ FA images were then merged into a single 4D image file (“all_FA”). The mean of all FA images was then created (“mean_FA”), which was fed into the FA skeletonisation program, creating a “mean_FA_skeleton”. Finally, all subjects’ FA data were projected onto the mean FA skeleton. This image was then thresholded (0.2), with the resulting binary skeleton mask defining the voxels used in all subsequent processing. A "distance map" was then created from the skeleton mask, which was used in the projection of FA onto the skeleton. Finally, the script took the 4D “all_FA” image (containing all subjects' aligned FA data) and, for each subject, the FA data were projected onto the mean FA skeleton. This resulted in a 4D image file containing the (projected) skeletonised FA data. The same pipeline was followed for AD and RD data. These files were then fed into GLM modelling for voxel-wise statistics. Second-level covariates included age and sex. TFCE (Smith and Nichols 2009) was used to correct for multiple comparisons (p-FWE<0.05; 5,000 permutations).

#### S-Methods 6: Conceptual replication of fixel-based analysis with Manual reconstruction of WM tracts of interest, using deterministic tractography

Using the ExploreDTI graphical toolbox(Leemans and Jones 2009; Leemans et al. 2009) (<http://www.exploredti.com/>; PROVIDI Lab, Utrecht, the Netherlands), we corrected for subject motion and eddy current distortions and reconstructed tract pathways using whole-brain deterministic WM tractography (maximal angle: 60°; step size: 1mm; minimal fiber length: 10mm; maximal fiber length: 500 mm; minimal FA for seed point selection: 0.1). Using manually drawn waypoint regions of interest on the direction-encoded FA maps in individual subject space, we generated three-dimensional reconstructions of each tract of interest (determined a priori, driven by our hypotheses on WM abnormalities in the hippocampal-diencephalic-cingulate network: fornix, dorsal and ventral cingulum, mammillothalamic tract, and the anterior thalamic radiation, especially since it comprises fibres from the principal anterior nuclei, as well as from the mediodorsal nuclei (Parent 1996); a posteriori, following whole-brain analyses, *supra*: anterior/superior corona radiata), based on protocols for each of those tracts (anterior-superior corona radiata, dorsal and ventral cingulum, anterior thalamic radiation (Wakana et al. 2007); mammillothalamic tract (Kamali et al. 2018); fornix (Metzler-Baddeley et al. 2011, 2013). Following tract reconstruction, we derived the median FA, AD, and RD values per tract per participant at each 1 mm step along each tract.

### Supplementary Figures

#### Figure S1

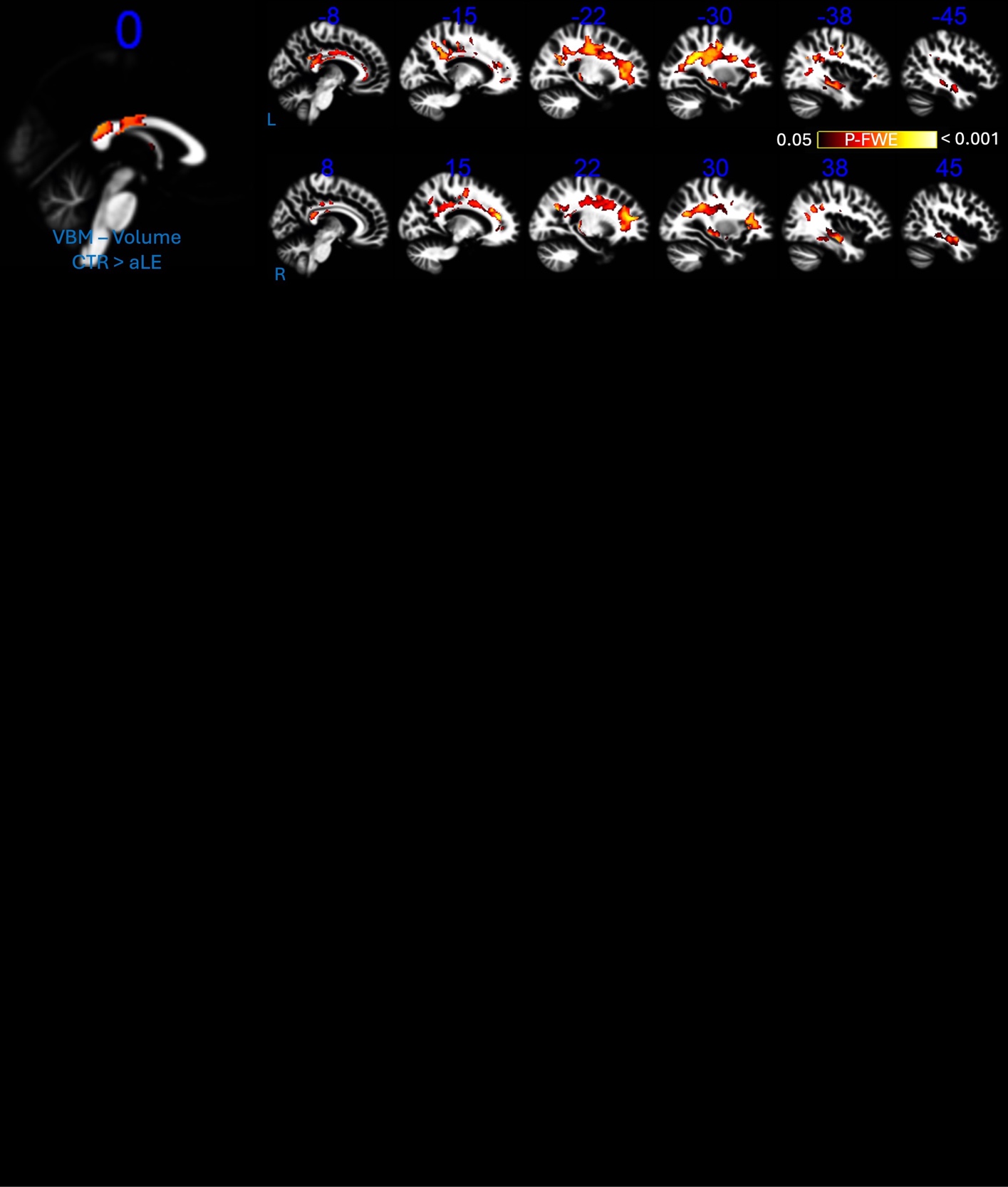

White-matter Voxel-Based Morphometry; **key:** aLE: autoimmune Limbic Encephalitis; CTR: healthy controls; VBM: Voxel-Based Morphometry; p-FWE: We used Threshold-Free Cluster Enhancement (TFCE) (Smith and Nichols 2009) to correct for multiple comparisons (p-FWE < 0.05; based on 5,000 permutations)

#### Figure S2

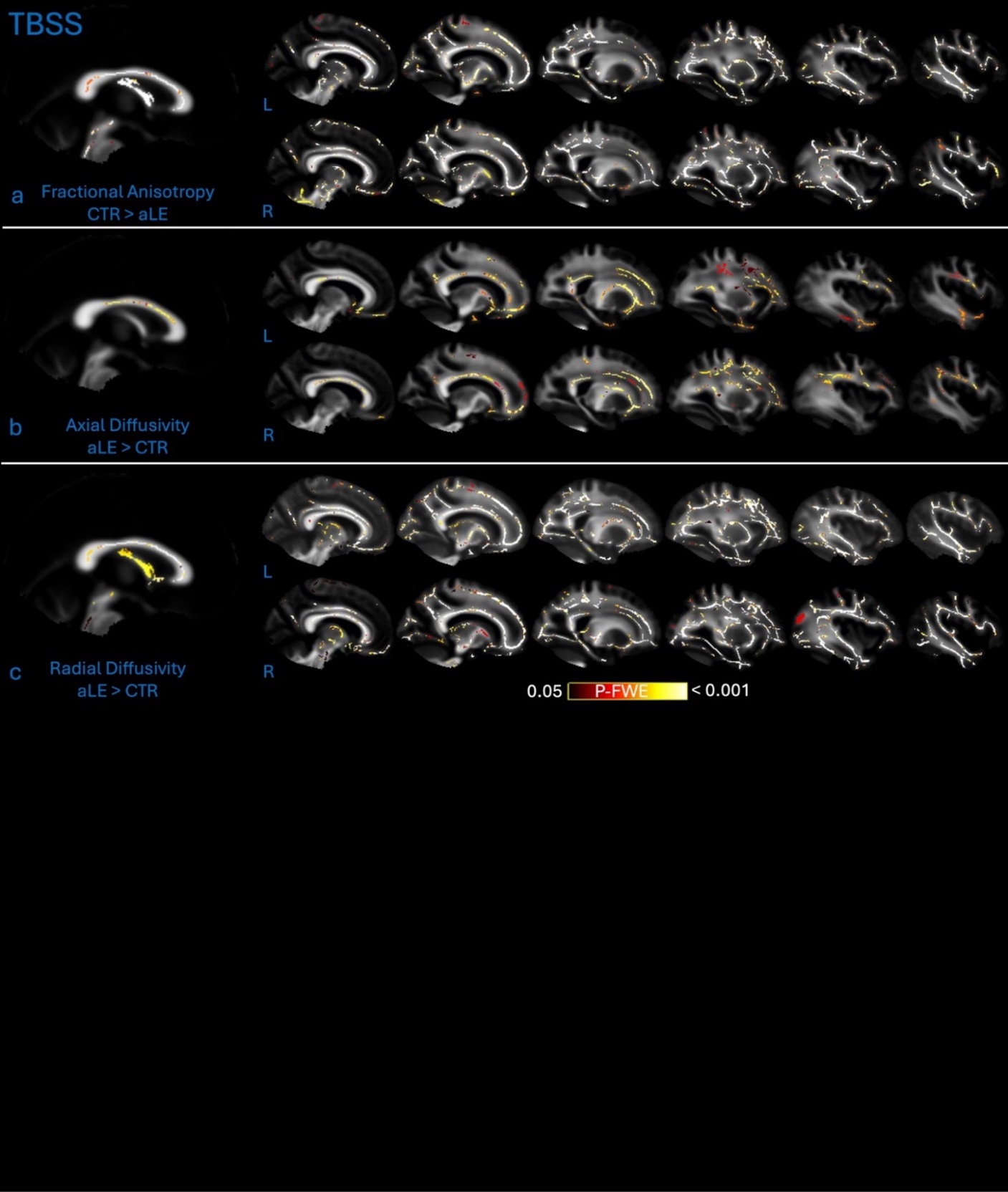

Tract-Based Spatial Statistics; **key:** aLE: autoimmune Limbic Encephalitis; CTR: healthy controls; L: left hemisphere; R: right hemisphere; We used Threshold-Free Cluster Enhancement (TFCE) (Smith and Nichols 2009) to correct for multiple comparisons (p-FWE < 0.05; based on 5,000 permutations)

#### Figure S3

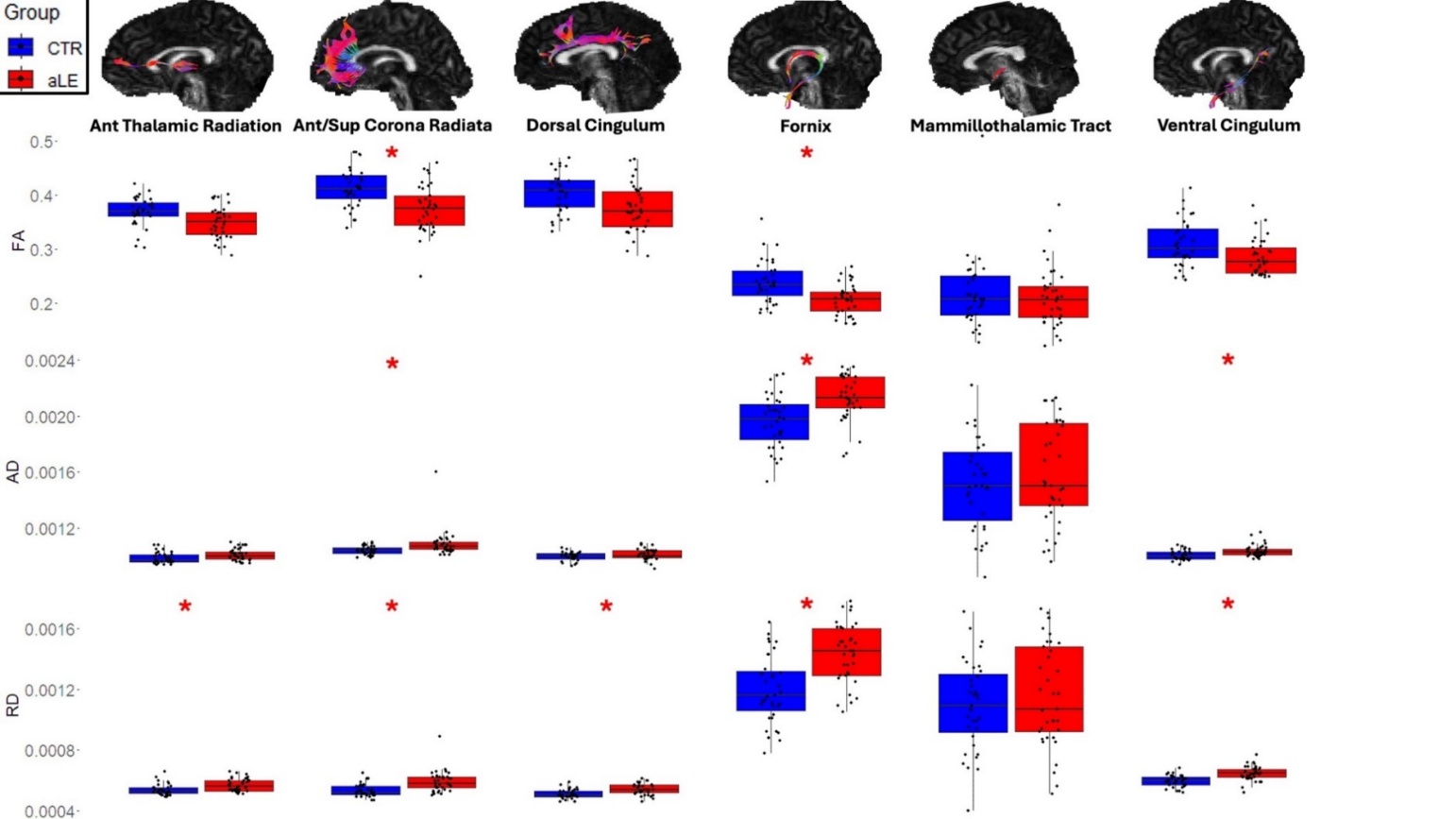

Median FA, AD, and RD, for manually reconstructed tracts using deterministic tractography: anterior thalamic radiation, anterior-superior corona radiata, dorsal cingulum, fornix, mammillothalamic tract, and ventral cingulum; **key:** AD: axial diffusivity; aLE: autoimmune Limbic Encephalitis; CTR: healthy controls; FA: fractional anisotropy; RD: radial diffusivity; *: survives correction for multiple testing using the Holm-Bonferroni procedure; line within each boxplot = median value; bottom of box = 25th %ile; top of box = 75th %ile; upper and lower whiskers = scores outside the middle 50; whiskers = 1.5 * interquartile range.

#### Figure S4

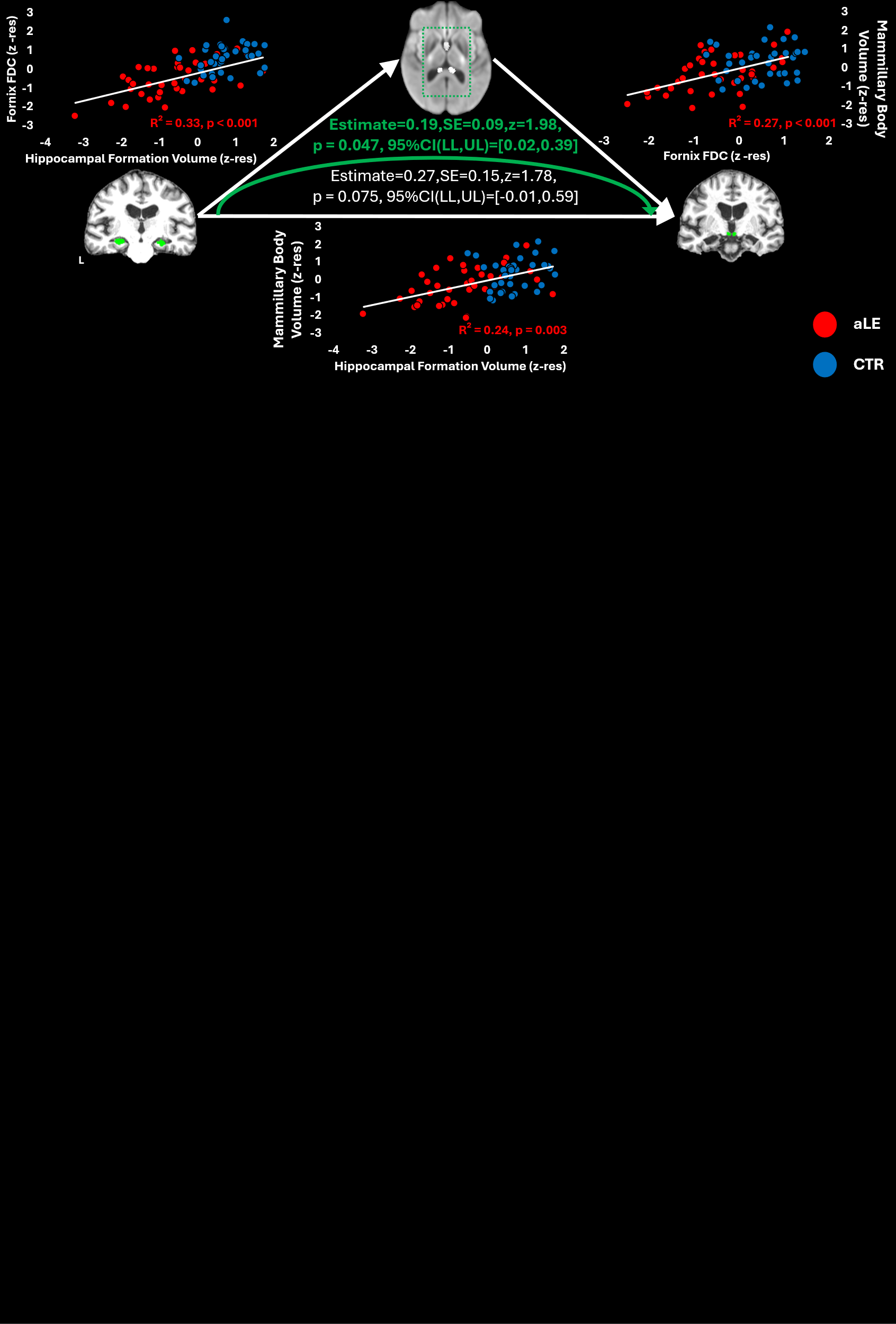

Mediation analysis among the total volume of the hippocampal formation (predictor; volume residualised against age, sex, intracranial volume, and scan source), fornical integrity (mediator; mean FDC, residualised against age and sex), and total mammillary bodies volume (outcome variable; volumes residualised against age, sex, total intracranial volume, and scan source); data from healthy controls are only illustrated for display purposes and are not used in the calculation of the R^2^ reported; **key:** aLE: autoimmune Limbic Encephalitis; CTR: healthy controls; FDC: Fiber Density and Cross-section; L: left hemisphere; R: right hemisphere; z-res: standardised residuals; SE: standard errors (bootstrapping with 5,000 samples); 95%CI(LL,UL) = 95% confidence intervals (lower limit,upper limit).

### Supplementary Tables

#### Table S1

| Structure | CTR | | aLE | | CTR vs. aLE | | | | |
| --- | --- | --- | --- | --- | --- | --- | --- | --- | --- |
|  | Mean  n vox | SD  n vox | Mean  n vox | SD  n vox | F | MSe | p-corr  (24) | η^2^_p_ | %  reduction |
| T H A L A M U S* | | | | | | | | | |
| Principal Anterior | 419.82 | 119.00 | 270.15 | 117.24 | 33.27 | 6694.68 | < 0.001 | 0.33 | 35.65 |
| Laterodorsal | 140.83 | 34.03 | 110.65 | 26.48 | 13.43 | 687.94 | 0.010 | 0.17 | 21.43 |
| Central Medial | 199.02 | 28.91 | 175.88 | 25.35 | 9.18 | 549.87 | 0.052 | 0.12 | 11.63 |
| Centrolateral | 126.41 | 22.68 | 110.72 | 22.49 | 6.48 | 472.15 | 0.105 | 0.09 | 12.41 |
| Centromedian | 555.21 | 81.88 | 467.36 | 114.64 | 9.66 | 8019.86 | 0.047*** | 0.13 | 15.82 |
| Limitans - Suprageniculate | 67.45 | 17.70 | 54.15 | 11.59 | 10.47 | 162.18 | 0.034 | 0.14 | 19.71 |
| Lateral Geniculate | 546.26 | 126.86 | 451.17 | 141.11 | 5.47 | 15035.34 | 0.112 | 0.08 | 17.41 |
| Lateral Posterior | 421.29 | 52.78 | 389.39 | 68.32 | 2.96 | 3059.49 | 0.179 | 0.04 | 7.57 |
| Mediodorsal lateral | 458.27 | 76.73 | 357.90 | 113.03 | 15.21 | 7214.64 | 0.005 | 0.19 | 21.90 |
| Mediodorsal medial | 1238.21 | 208.43 | 961.69 | 276.19 | 19.61 | 40058.12 | < 0.001 | 0.23 | 22.33 |
| Medial Geniculate | 467.72 | 93.17 | 401.81 | 107.13 | 4.70 | 8964.22 | 0.135 | 0.07 | 14.09 |
| Medial Ventral - Reuniens | 53.64 | 25.90 | 36.32 | 15.64 | 8.30 | 308.33 | 0.063 | 0.11 | 32.28 |
| Parafascicular | 111.79 | 25.72 | 90.63 | 30.49 | 6.37 | 540.09 | 0.105 | 0.09 | 18.93 |
| Pulvinar - Anterior | 524.30 | 65.74 | 462.78 | 88.76 | 7.82 | 4194.33 | 0.065 | 0.10 | 11.73 |
| Pulvinar - Inferior | 426.95 | 43.26 | 377.05 | 75.56 | 7.91 | 3187.79 | 0.065 | 0.11 | 11.69 |
| Pulvinar Lateral | 340.70 | 39.00 | 297.55 | 52.13 | 11.61 | 1740.79 | 0.021 | 0.15 | 12.67 |
| Pulvinar Medial (lateral segment) | 1501.14 | 152.15 | 1340.27 | 233.77 | 8.48 | 31948.45 | 0.063 | 0.11 | 10.72 |
| Pulvinar Medial (medial segment) | 549.88 | 124.94 | 379.91 | 129.49 | 31.39 | 9989.85 | < 0.001 | 0.32 | 30.91 |
| Ventral Anterior | 764.24 | 162.35 | 605.42 | 215.02 | 8.44 | 26437.13 | 0.063 | 0.11 | 20.78 |
| Ventral Anterior (magnocellular) | 42.86 | 6.92 | 41.07 | 7.02 | 0.40 | 41.98 | 0.531 | 0.01 | 4.17 |
| Ventrolateral anterior | 794.70 | 160.47 | 656.69 | 200.94 | 6.44 | 25634.25 | 0.105 | 0.09 | 17.37 |
| Ventrolateral posterior | 1299.96 | 252.56 | 1075.10 | 293.66 | 8.78 | 55805.89 | 0.059 | 0.12 | 17.30 |
| Ventral Posterolateral | 1474.60 | 218.28 | 1352.12 | 205.78 | 3.77 | 36128.11 | 0.170 | 0.05 | 8.31 |
| HYPOTHALAMUS** | | | | | | | | | |
| Mammillary Bodies | 106.18 | 14.35 | 98.87 | 14.25 | 9.24 | 172.02 | 0.048 | 0.09 | 6.89 |

Comparisons between aLE patients and CTRs on the volumes of thalamic nuclei (estimated by ‘FreeSurfer-CNN’(Tregidgo et al. 2023)) and the mammillary bodies (estimated by FreeSurfer-ScLimbic) (Greve et al. 2021); **key:** p-corr: p value corrected for multiple comparisons, using the Holm-Bonferroni sequential correction method (Holm 1979); CTR: healthy controls; aLE: autoimmune Limbic Encephalitis; *n* vox: number of voxels; *: Comparisons between CTRs (n=35) and aLE patients (n = 37), i.e., participants for whom both T_1_-weighted structural MRIs and diffusion data were available, on thalamic volumes estimated by FreeSurfer-CNN (ANCOVAs included age, sex, and TIV as covariates); **: Comparisons on mammillary body volumes estimated by ScLimbic, between CTRs (n=67) and aLE patients (n=38); ANCOVAs included age, sex, total intracranial volume, and scan source as covariates; ***: as the data for this volume were not normally distributed, we iterated this comparison using Quade’s non-parametric ANCOVA; unlike other volumes that were not normally distributed, the p-corr value for this ANCOVA did not approach significance (p-corr(24) = 0.104; voxel size: 1 × 1 × 1 mm).

#### Table S2

| Structure | CTR (n=67) | | aLE (n=38) | | CTR vs aLE | | | | |
| --- | --- | --- | --- | --- | --- | --- | --- | --- | --- |
|  | Mean  n vox | SD  n vox | Mean  n vox | SD  n vox | *F*_(1,99)_ | MSe | p-corr(16) | η^2^_p_ | % reduction |
| THALAMUS | | | | | | | | | |
| Medial Geniculate | 114.88 | 16.75 | 108.47 | 17.60 | 6.21 | 178.97 | 0.144 | 0.06 | 5.58 |
| Centromedian | 207.73 | 39.27 | 185.87 | 35.74 | 7.68 | 947.32 | 0.080 | 0.07 | 10.52 |
| Mediodorsal-Parafascicular | 1204.51 | 174.72 | 1079.95 | 186.30 | 20.91 | 18887.83 | < 0.001 | 0.17 | 10.34 |
| Habenula | 47.28 | 9.35 | 46.03 | 10.94 | 1.22 | 100.39 | 0.545 | 0.01 | 2.66 |
| Anteroventral | 196.24 | 58.57 | 152.13 | 50.50 | 31.94 | 1736.60 | < 0.001 | 0.24 | 22.48 |
| Ventral Anterior | 542.52 | 88.38 | 519.82 | 65.07 | 4.07 | 3766.19 | 0.372 | 0.04 | 4.19 |
| Ventrolateral Anterior | 157.97 | 21.75 | 153.74 | 20.70 | 3.33 | 279.03 | 0.378 | 0.03 | 2.68 |
| Ventrolateral Posterior | 1598.93 | 243.42 | 1514.34 | 221.15 | 6.13 | 36879.40 | 0.144 | 0.06 | 5.29 |
| Ventral Posterolateral | 563.24 | 68.34 | 547.61 | 68.59 | 2.93 | 3409.76 | 0.378 | 0.03 | 2.78 |
| Pulvinar | 2368.52 | 344.15 | 2135.18 | 334.86 | 22.88 | 59108.26 | < 0.001 | 0.19 | 9.85 |
| Lateral Geniculate | 197.15 | 36.31 | 177.26 | 37.09 | 6.39 | 1061.87 | 0.144 | 0.06 | 10.09 |
| HYPOTHALAMUS | | | | | | | | | |
| Anterior-Inferior | 25.36 | 8.16 | 22.67 | 6.80 | 0.03 | 56.45 | 0.864 | < 0.01 | 10.62 |
| Anterior-Superior | 37.75 | 7.64 | 32.59 | 8.63 | 3.24 | 57.17 | 0.378 | 0.03 | 13.67 |
| Posterior | 221.40 | 26.35 | 198.99 | 32.73 | 9.70 | 760.31 | 0.031 | 0.09 | 10.12 |
| Tubular-Inferior | 258.63 | 27.86 | 247.12 | 36.51 | 3.87 | 961.12 | 0.372 | 0.04 | 4.45 |
| Tubular-Superior | 212.24 | 25.52 | 200.50 | 28.83 | 3.54 | 643.75 | 0.378 | 0.03 | 5.53 |

Comparisons between CTRs (n=67) and aLE patients (n = 38), i.e., all participants with available T_1_-weighted structural MRI data, on volumes of thalamic nuclei, estimated by HIPS-THOMAS (Vidal et al. 2024), and hypothalamic volumes, estimated by the HypothalamicSubunits tool (Billot et al. 2020). Univariate ANCOVAs involved age at scanning, sex, total intracranial volume and scan source as covariates; **key:** p-corr: p value corrected for multiple comparisons, using the Holm-Bonferroni sequential correction method; Hem: Hemisphere; L, R: Left, Right hemisphere; CTR: healthy controls; aLE: autoimmune Limbic Encephalitis; n vox: number of voxels (voxel size: 1 × 1 × 1 mm).

#### Table S3

| Memory Domain (Outcome) | Diencephalic Volume (Mediator) | Diencephalic volume - memory (bivariate correlation) | | Hippocampal/Subicular (Subfield) Volume (Predictor) | Predictor-Outcome relationship (bivariate correlation) | | Mediator-Predictor relationship (bivariate correlation) | | **Mediator-Outcome relationship, controlling for Predictor (partial correlation)** | | **Mediation** | | | | | | | |
| --- | --- | --- | --- | --- | --- | --- | --- | --- | --- | --- | --- | --- | --- | --- | --- | --- | --- | --- |
|  |  |  |  |  |  |  |  |  |  |  | Direct Effect | | | | **Indirect Effect** | | | |
|  |  | r | p* | (max r) | r | p | r | p | **r** | **p** | Estimate | SE | z | CI95% LL-UL | **Estimate** | **SE** | **z** | **CI95% LL-UL** |
| Anterograde Retrieval | Mammillary  Body | 0.50 | 0.001 | Presubiculum (head) | 0.41 | 0.011 | 0.34 | 0.038 | **0.42** | **0.010** | 0.20 | 0.11 | 1.76 | -0.02,0.43 | **0.10** | **0.07** | **1.50** | **0.01,0.27** |
|  |  |  |  | Manually delineated Hippocampus | 0.32 | 0.052 | 0.47 | 0.003 | **0.42** | **0.010** | 0.09 | 0.12 | 0.72 | -0.12,0.35 | **0.18** | **0.09** | **2.04** | **0.04,0.38** |
|  | Laterodorsal  Nucleus | 0.46 | 0.004 | Presubiculum (head) | 0.41 | 0.011 | 0.59 | < 0.001 | **0.31** | **0.067** | 0.14 | 0.15 | 0.95 | -0.16,0.41 | **0.16** | **0.08** | **1.85** | **0.01,0.34** |
|  |  |  |  | Manually delineated Hippocampus | 0.32 | 0.052 | 0.49 | 0.002 | **0.38** | **0.024** | 0.09 | 0.15 | 0.62 | -0.22,0.36 | **0.17** | **0.08** | **2.07** | **0.05,0.37** |
| Remote  Autobiographical | Medial (m)  Pulvinar | 0.62 | < 0.001 | Subiculum (body) | 0.53 | 0.001 | 0.58 | < 0.001 | **0.47** | **0.006** | 0.84 | 0.63 | 1.32 | -0.32,2.18 | **1.07** | **0.42** | **2.58** | **0.36,1.96** |
|  |  |  |  | Manually delineated Hippocampus | 0.42 | 0.013 | 0.52 | < 0.001 | **0.53** | **0.002** | 0.63 | 0.71 | 0.89 | -0.74,2.04 | **1.14** | **0.45** | **2.52** | **0.43,2.15** |
| Verbal  Recognition | Mammillary  Body | 0.54 | < 0.001 | Subiculum (head) | 0.36 | 0.028 | 0.45 | 0.005 | **0.46** | **0.005** | 0.14 | 0.17 | 0.83 | -0.17,0.49 | **0.17** | **0.09** | **2.00** | **0.04,0.36** |
|  |  |  |  | Manually delineated Hippocampus | 0.30 | 0.072 | 0.47 | 0.003 | **0.47** | **0.004** | 0.05 | 0.17 | 0.28 | -0.23,0.44 | **0.22** | **0.09** | **2.44** | **0.07,0.42** |
| Visual  Recognition | Laterodorsal  Nucleus | 0.58 | < 0.001 | CA4 (head) | 0.45 | 0.005 | 0.60 | 0.001 | **0.45** | **0.008** | 0.13 | 0.18 | 0.71 | -0.23,0.47 | **0.32** | **0.11** | **2.95** | **0.12,0.56** |
|  |  |  |  | Manually delineated Hippocampus | 0.29 | 0.088 | 0.49 | 0.002 | **0.54** | **0.001** | -0.01 | 0.17 | -0.04 | -0.38,0.30 | **0.32** | **0.09** | **3.52** | **0.17,0.53** |

Mediation analyses on memory scores (Argyropoulos et al. 2019), volumes of diencephalic segmentations, and volumes of hippocampal segmentations (total hippocampal volume, manually delineated (Argyropoulos et al. 2019), or volume of automatically segmented hippocampal/subicular subfields with highest Pearson coefficient value for each memory score, deliberately examined at uncorrected levels) ; **key: p***: these relationships survive correction for multiple correlations using the Holm-Bonferroni procedure (Holm 1979) (see main text); L, R: left, right hemisphere; r: Pearson’s correlation coefficient; SE: standard errors (bootstrapping with 5,000 samples); CI95%LL-UL: 95% confidence intervals (lower limit-upper limit); FDC: fiber density and cross-section.

#### Table S4

| Tracts | CTR (n=35) | | aLE (n=37) | | CTR vs aLE | | | |
| --- | --- | --- | --- | --- | --- | --- | --- | --- |
|  | Mean | SD | Mean | SD | *F*_(1,68)_ | MSe | η^2^_p_ | p-corr (33) |
| Anterior Commissure | 0.41 | 0.06 | 0.35 | 0.06 | 17.83 | 0.002 | 0.21 | 0.002 |
| Anterior Thalamic Radiation | 0.38 | 0.05 | 0.34 | 0.04 | 7.79 | 0.001 | 0.10 | 0.055 |
| Arcuate Fasciculus | 0.39 | 0.05 | 0.35 | 0.05 | 12.12 | 0.001 | 0.15 | 0.016 |
| Cingulum | 0.35 | 0.04 | 0.31 | 0.04 | 12.17 | 0.001 | 0.15 | 0.016 |
| Corpus Callosum | 0.44 | 0.05 | 0.39 | 0.05 | 11.67 | 0.002 | 0.15 | 0.018 |
| Cortico-Spinal Tract | 0.63 | 0.08 | 0.55 | 0.08 | 12.42 | 0.004 | 0.15 | 0.015 |
| Fornix | 0.40 | 0.06 | 0.32 | 0.07 | 32.64 | 0.002 | 0.32 | < 0.001 |
| Fronto-Pontine Tract | 0.50 | 0.06 | 0.46 | 0.05 | 11.16 | 0.002 | 0.14 | 0.022 |
| Inferior Longitudinal Fasciculus | 0.44 | 0.05 | 0.39 | 0.04 | 17.13 | 0.002 | 0.20 | 0.003 |
| Inferior Occipito-Frontal Fasciculus | 0.44 | 0.05 | 0.39 | 0.05 | 16.66 | 0.001 | 0.20 | 0.003 |
| Middle Cerebellar Peduncle | 0.50 | 0.07 | 0.46 | 0.07 | 4.16 | 0.005 | 0.06 | 0.136 |
| Middle Longitudinal Fasciculus | 0.41 | 0.04 | 0.37 | 0.04 | 18.19 | 0.001 | 0.21 | 0.002 |
| Optic Radiation | 0.44 | 0.05 | 0.39 | 0.05 | 13.96 | 0.002 | 0.17 | 0.008 |
| Parieto-Occipital Pontine | 0.51 | 0.05 | 0.46 | 0.05 | 15.60 | 0.002 | 0.19 | 0.005 |
| Striato-Fronto-Orbital | 0.40 | 0.05 | 0.37 | 0.05 | 4.14 | 0.001 | 0.06 | 0.136 |
| Striato-Occipital | 0.46 | 0.06 | 0.41 | 0.05 | 16.48 | 0.002 | 0.20 | 0.003 |
| Striato-Parietal | 0.45 | 0.04 | 0.40 | 0.05 | 15.74 | 0.001 | 0.19 | 0.005 |
| Striato-Post-Central | 0.47 | 0.05 | 0.42 | 0.05 | 9.00 | 0.002 | 0.12 | 0.041 |
| Striato-Pre-Central | 0.50 | 0.06 | 0.45 | 0.06 | 8.85 | 0.003 | 0.12 | 0.041 |
| Striato-Prefrontal | 0.41 | 0.05 | 0.37 | 0.05 | 6.49 | 0.001 | 0.09 | 0.079 |
| Striato-Premotor | 0.42 | 0.06 | 0.39 | 0.05 | 3.72 | 0.002 | 0.05 | 0.136 |
| Superior Cerebellar Peduncle | 0.49 | 0.05 | 0.42 | 0.05 | 31.41 | 0.002 | 0.32 | < 0.001 |
| Superior Longitudinal Fasciculus I | 0.38 | 0.04 | 0.35 | 0.04 | 6.45 | 0.001 | 0.09 | 0.079 |
| Superior Longitudinal Fasciculus II | 0.38 | 0.05 | 0.34 | 0.05 | 10.61 | 0.002 | 0.14 | 0.026 |
| Superior Longitudinal Fasciculus III | 0.39 | 0.05 | 0.34 | 0.05 | 9.79 | 0.002 | 0.13 | 0.034 |
| Superior Thalamic Radiation | 0.57 | 0.07 | 0.51 | 0.07 | 8.32 | 0.004 | 0.11 | 0.047 |
| Thalamo-Occipital | 0.44 | 0.05 | 0.39 | 0.05 | 14.78 | 0.002 | 0.18 | 0.006 |
| Thalamo-Parietal | 0.45 | 0.04 | 0.40 | 0.05 | 15.50 | 0.001 | 0.19 | 0.005 |
| Thalamo-Post-Central | 0.52 | 0.06 | 0.46 | 0.06 | 10.10 | 0.003 | 0.13 | 0.031 |
| Thalamo-Pre-Central | 0.52 | 0.06 | 0.47 | 0.06 | 9.79 | 0.003 | 0.13 | 0.034 |
| Thalamo-Prefrontal | 0.41 | 0.05 | 0.37 | 0.04 | 7.31 | 0.001 | 0.10 | 0.061 |
| Thalamo-Premotor | 0.42 | 0.05 | 0.39 | 0.05 | 5.21 | 0.002 | 0.07 | 0.102 |
| Uncinate Fasciculus | 0.43 | 0.05 | 0.38 | 0.05 | 14.59 | 0.002 | 0.18 | 0.006 |

Mean FDC values per tract, collapsing across hemisphere for tracts automatically delineated separately for left and right hemisphere by TractSeg. Univariate ANCOVAs involved age at scanning and sex as between subjects covariates of no interest. **key:** aLE: autoimmune Limbic Encephalitis; CTR: healthy controls; FDC: Fiber Density and Cross-section; p-corr: Bonferroni-Holm correction for multiple testing.

#### Table S5

| Full Model | | | | | | |
| --- | --- | --- | --- | --- | --- | --- |
| Effects and Interactions | Sum Sq | Mean Sq | Df1 | Df2 | F | p |
| FX | 1.70 | 1.70 | 1 | 22.76 | 2.94 | 0.100 |
| Material Type | 23.64 | 11.82 | 2 | 159.66 | 20.46 | 0.000 |
| Paradigm | 0.30 | 0.30 | 1 | 159.66 | 0.53 | 0.469 |
| Process | 0.01 | 0.01 | 1 | 159.66 | 0.01 | 0.906 |
| FX*Material Type | 2.72 | 1.36 | 2 | 159.66 | 2.36 | 0.098 |
| FX*Paradigm | 0.18 | 0.18 | 1 | 159.66 | 0.32 | 0.575 |
| Material Type*Paradigm | 1.78 | 0.89 | 2 | 159.66 | 1.54 | 0.218 |
| FX*Process | 2.93 | 2.93 | 1 | 159.66 | 5.08 | 0.026 |
| Material Type*Process | 2.32 | 1.16 | 2 | 159.66 | 2.01 | 0.138 |
| Paradigm*Process | 4.12 | 4.12 | 1 | 159.66 | 7.14 | 0.008 |
| FX*Material Type*Paradigm | 0.87 | 0.44 | 2 | 159.66 | 0.76 | 0.471 |
| FX*Material Type*Process | 9.01 | 4.50 | 2 | 159.66 | 7.80 | 0.001 |
| FX*Paradigm*Process | 5.26 | 5.26 | 1 | 159.66 | 9.11 | 0.003 |
| Material Type*Paradigm*Process | 1.61 | 0.80 | 2 | 159.66 | 1.39 | 0.252 |
| FX*Material Type*Paradigm*Process | 5.32 | 2.66 | 2 | 159.66 | 4.60 | 0.011 |
| Separate Models for different Process and Material types | | | | | | |
| Effect of FX on… | Sum Sq | Mean Sq | Df1 | Df2 | F | p-corr(6) |
| Familiarity – Faces | 0.00 | 0.00 | 1 | 27.97 | 0.02 | <0.999 |
| Familiarity – Scenes | 10.78 | 10.78 | 1 | 7.91 | 16.31 | 0.023 |
| Familiarity – Words | 0.29 | 0.29 | 1 | 26.12 | 1.12 | <0.999 |
| Recollection – Faces | 0.76 | 0.76 | 1 | 16.53 | 2.95 | 0.523 |
| Recollection – Scenes | 0.17 | 0.17 | 1 | 19.07 | 0.37 | <0.999 |
| Recollection - Words | 0.00 | 0.00 | 1 | 21.18 | 0.01 | <0.999 |
| Separate Models for different Process types | | | | | | |
| Effect of FX on… | | | Estimate | Std. Error | t value | p-corr(2) |
| Recollection | | | 0.30 | 0.20 | 1.44 | 0.199 |
| Familiarity | | | 0.64 | 0.20 | 3.13 | 0.041 |

Effects of FX (and interactions with other variables) on familiarity/recollection estimates (using linear mixed effects models - “lmer” in R) for the aLE patients who had completed our custom-made behavioural tasks dissociating recollection from familiarity (Argyropoulos et al. 2022); we used fully factorial fixed effects of mean FDC, Process (Familiarity, Recollection), Material-Type (Faces, Scenes, Words), Paradigm (the two tasks involved), and Hemisphere (left, right). A single random intercept across participants was used. The results were reported in terms of Type III ANOVA using Satterthwaite’s method for adjusting degrees of freedom. We then iterated this analysis, separately for Process and Process-and-Material-Type; **key:** FX: fornix integrity (mean FDC, residualised against age and sex); p-corr: p values corrected according to the number of models the same analyses were iterated for, using the Holm-Bonferroni method (Holm 1979).

#### Table S6

| Structure | CTR | | LGI-1 aLE (n=14) | | CTR vs. LGI-1 aLE | | | | |
| --- | --- | --- | --- | --- | --- | --- | --- | --- | --- |
|  | Mean  n vox | SD  n vox | Mean  n vox | SD  n vox | F | MSe | p-corr (24) | η^2^_p_ | % reduction |
| T H A L A M U S* | | | | | | | | | |
| Principal Anterior | 419.82 | 119.00 | 237.50 | 91.70 | 23.72 | 5924.87 | < 0.001 | 0.35 | 43.43 |
| Laterodorsal | 140.83 | 34.03 | 111.06 | 22.47 | 5.35 | 703.84 | 0.238 | 0.11 | 21.14 |
| Central Medial | 199.02 | 28.91 | 173.76 | 15.47 | 7.20 | 397.63 | 0.133 | 0.14 | 12.69 |
| Centrolateral | 126.41 | 22.68 | 113.20 | 21.08 | 3.53 | 403.98 | 0.302 | 0.07 | 10.45 |
| Centromedian | 555.21 | 81.88 | 466.92 | 108.65 | 7.71 | 5312.48 | 0.113 | 0.15 | 15.90 |
| Limitans - Suprageniculate | 67.45 | 17.70 | 52.00 | 8.07 | 5.48 | 174.88 | 0.238 | 0.11 | 22.90 |
| Lateral Geniculate | 546.26 | 126.86 | 442.00 | 129.30 | 3.35 | 13155.76 | 0.302 | 0.07 | 19.09 |
| Lateral Posterior | 421.29 | 52.78 | 389.89 | 51.54 | 4.16 | 1947.31 | 0.293 | 0.09 | 7.45 |
| Mediodorsal lateral | 458.27 | 76.73 | 345.00 | 102.60 | 21.37 | 4199.39 | 0.001 | 0.33 | 24.72 |
| Mediodorsal medial | 1238.21 | 208.43 | 924.05 | 233.89 | 25.47 | 24646.27 | < 0.001 | 0.37 | 25.37 |
| Medial Geniculate | 467.72 | 93.17 | 382.18 | 78.15 | 5.85 | 6891.15 | 0.218 | 0.12 | 18.29 |
| Medial Ventral - Reuniens | 53.64 | 25.90 | 32.28 | 10.78 | 4.39 | 380.08 | 0.293 | 0.09 | 39.82 |
| Parafascicular | 111.79 | 25.72 | 86.40 | 24.61 | 4.89 | 363.57 | 0.258 | 0.10 | 22.71 |
| Pulvinar - Anterior | 524.30 | 65.74 | 456.16 | 89.80 | 9.13 | 3051.03 | 0.067 | 0.17 | 13.00 |
| Pulvinar - Inferior | 426.95 | 43.26 | 366.58 | 62.94 | 9.70 | 2090.56 | 0.055 | 0.18 | 14.14 |
| Pulvinar Lateral | 340.70 | 39.00 | 295.15 | 36.79 | 12.49 | 1078.22 | 0.019 | 0.22 | 13.37 |
| Pulvinar Medial (lateral segment) | 1501.14 | 152.15 | 1316.50 | 192.96 | 12.51 | 19961.15 | 0.019 | 0.22 | 12.30 |
| Pulvinar Medial (medial segment) | 549.88 | 124.94 | 351.10 | 100.47 | 23.65 | 8699.87 | 0.000 | 0.35 | 36.15 |
| Ventral Anterior | 764.24 | 162.35 | 583.67 | 206.42 | 8.32 | 19954.88 | 0.091 | 0.16 | 23.63 |
| Ventral Anterior (magnocellular) | 42.86 | 6.92 | 43.32 | 7.60 | 0.29 | 43.05 | 0.595 | 0.01 | -1.07 |
| Ventrolateral anterior | 794.70 | 160.47 | 638.40 | 180.87 | 6.42 | 19460.94 | 0.179 | 0.13 | 19.67 |
| Ventrolateral posterior | 1299.96 | 252.56 | 1051.33 | 290.36 | 10.72 | 42941.79 | 0.037 | 0.20 | 19.13 |
| Ventral Posterolateral | 1474.60 | 218.28 | 1383.82 | 174.41 | 1.44 | 29167.93 | 0.472 | 0.03 | 6.16 |
| HYPOTHALAMUS** | | | | | | | | | |
| Mammillary Bodies | 106.18 | 14.35 | 102.29 | 13.77 | 3.63 | 165.51 | 0.302 | 0.05 | 3.67 |

Comparisons between LGI-1 aLE patients and CTRs on the volumes of thalamic nuclei estimated by ‘FreeSurfer-CNN’ (Tregidgo et al. 2023) and the mammillary bodies, estimated by FreeSurfer-ScLimbic (Greve et al. 2021); **key:** p-corr: p value corrected for multiple comparisons, using the Holm-Bonferroni sequential correction method; CTR: healthy controls; LGI-1 aLE: anti-leucine-rich glioma inactivated autoimmune limbic encephalitis; *n* vox: number of voxels; *: Comparisons between CTRs (n=35) and LGI-1 aLE patients (n = 14), i.e., participants for whom both T_1_-weighted structural MRIs and diffusion data were available, on thalamic volumes estimated by FreeSurfer-CNN (ANCOVAs included age, sex, and TIV as covariates); Comparisons on mammillary body volumes estimated by ScLimbic, between CTRs (n=67) and LGI-1 aLE patients (n=14); ANCOVAs included age, sex, total intracranial volume, and scan source as covariates; voxel size: 1 × 1 × 1 mm).

#### Table S7

| Structure | CTR (n=67) | | LGI-1 aLE (n=14) | | CTR vs LGI-1 aLE | | | | |
| --- | --- | --- | --- | --- | --- | --- | --- | --- | --- |
|  | Mean  n vox | SD  n vox | Mean  n vox | SD  n vox | *F*_(1,75)_ | MSe | η^2^_p_ | p-corr(16) | % reduction |
| THALAMUS | | | | | | | | | |
| Medial Geniculate | 114.88 | 16.75 | 102.29 | 13.40 | 9.23 | 155.83 | 0.11 | 0.043 | 10.96 |
| Centromedian | 207.73 | 39.27 | 174.64 | 34.37 | 7.42 | 903.70 | 0.09 | 0.096 | 15.93 |
| Mediodorsal-Parafascicular | 1204.51 | 174.72 | 1057.71 | 175.76 | 17.11 | 17318.48 | 0.19 | 0.001 | 12.19 |
| Habenula | 47.28 | 9.35 | 44.71 | 12.43 | 1.42 | 100.71 | 0.02 | > 0.999 | 5.43 |
| Anteroventral | 196.24 | 58.57 | 145.36 | 57.04 | 17.86 | 1699.73 | 0.19 | 0.001 | 25.93 |
| Ventral Anterior | 542.52 | 88.38 | 521.64 | 64.24 | 2.24 | 3806.86 | 0.03 | > 0.999 | 3.85 |
| Ventrolateral Anterior | 157.97 | 21.75 | 157.50 | 18.33 | 0.77 | 270.49 | 0.01 | > 0.999 | 0.30 |
| Ventrolateral Posterior | 1598.93 | 243.42 | 1507.29 | 222.00 | 3.65 | 38033.07 | 0.05 | 0.660 | 5.73 |
| Ventral Posterolateral | 563.24 | 68.34 | 545.93 | 70.18 | 1.31 | 3229.52 | 0.02 | > 0.999 | 3.07 |
| Pulvinar | 2368.52 | 344.15 | 2113.36 | 279.77 | 13.49 | 53665.49 | 0.15 | 0.006 | 10.77 |
| Lateral Geniculate | 197.15 | 36.31 | 173.36 | 31.35 | 3.57 | 994.00 | 0.05 | 0.660 | 12.07 |
| HYPOTHALAMUS | | | | | | | | | |
| Anterior-Inferior | 25.36 | 8.16 | 24.93 | 7.27 | 0.60 | 61.50 | 0.01 | > 0.999 | 1.70 |
| Anterior-Superior | 37.75 | 7.64 | 34.79 | 7.20 | 0.01 | 51.29 | 0.00 | > 0.999 | 7.84 |
| Posterior | 221.40 | 26.35 | 200.58 | 35.92 | 3.06 | 675.91 | 0.04 | 0.759 | 9.40 |
| Tubular-Inferior | 258.63 | 27.86 | 254.72 | 42.16 | 0.84 | 918.28 | 0.01 | > 0.999 | 1.51 |
| Tubular-Superior | 212.24 | 25.52 | 203.89 | 25.16 | 2.21 | 564.80 | 0.03 | > 0.999 | 3.94 |

Comparisons between CTRs (n=67) and LGI-1 aLE patients (n = 14) on volumes of thalamic nuclei, estimated by HIPS-THOMAS (Vidal et al. 2024), and hypothalamic volumes, estimated by the HypothalamicSubunits tool (Billot et al. 2020). Univariate ANCOVAs involved age at scanning, sex, total intracranial volume and scan source as covariates. Comparisons between this subgroup and the rest (n=24) of the patient cohort would not be informative, given that: i) 10 patients that were reported as seronegative may have had LGI-1 aLE, despite that anti-LGI-1 was not detectable in routine clinical practice at the time of screening; ii) only 2 patients were positive for anti-glutamic acid decarboxylase autoantibody, iii) 1 presented with autoantibodies characteristic of paraneoplastic encephalitis, iv) 4 presented with dual seropositive antibodies of contactin-associated protein-like 2 and LGI-1, and v) 7 were found positive for antibodies targeting the voltage-gated potassium channel complex, although there was no further information on the particular antibodies (Table 2 in (Argyropoulos et al. 2019)); **key:** p-corr: p value corrected for multiple comparisons, using the Holm-Bonferroni sequential correction method; Hem: Hemisphere; L, R: Left, Right hemisphere; CTR: healthy controls; LGI-1 aLE: anti-leucine-rich glioma inactivated autoimmune limbic encephalitis; n vox: number of voxels (voxel size: 1 × 1 × 1 mm).

#### Table S8

| Tracts | CTR (n=35) | | LGI-1 aLE (n=14) | | CTR vs LGI-1 aLE | | | |
| --- | --- | --- | --- | --- | --- | --- | --- | --- |
|  | Mean | SD | Mean | SD | *F*_(1,45)_ | MSe | η^2^_p_ | p-corr (33) |
| Anterior Commissure | 0.41 | 0.06 | 0.35 | 0.06 | 5.76 | 0.002 | 0.11 | 0.247 |
| Anterior Thalamic Radiation | 0.38 | 0.05 | 0.34 | 0.05 | 3.65 | 0.002 | 0.08 | 0.374 |
| Arcuate Fasciculus | 0.39 | 0.05 | 0.35 | 0.05 | 5.51 | 0.001 | 0.11 | 0.256 |
| Cingulum | 0.35 | 0.04 | 0.31 | 0.05 | 5.04 | 0.001 | 0.10 | 0.277 |
| Corpus Callosum | 0.44 | 0.05 | 0.38 | 0.06 | 6.25 | 0.002 | 0.12 | 0.241 |
| Cortico-Spinal Tract | 0.63 | 0.08 | 0.54 | 0.06 | 9.29 | 0.004 | 0.17 | 0.095 |
| Fornix | 0.40 | 0.06 | 0.32 | 0.06 | 13.80 | 0.002 | 0.23 | 0.018 |
| Fronto-Pontine Tract | 0.50 | 0.06 | 0.45 | 0.04 | 8.45 | 0.002 | 0.16 | 0.130 |
| Inferior Longitudinal Fasciculus | 0.44 | 0.05 | 0.39 | 0.04 | 7.54 | 0.002 | 0.14 | 0.165 |
| Inferior Occipito-Frontal Fasciculus | 0.44 | 0.05 | 0.38 | 0.05 | 9.32 | 0.002 | 0.17 | 0.095 |
| Middle Cerebellar Peduncle | 0.50 | 0.07 | 0.45 | 0.07 | 2.09 | 0.005 | 0.04 | 0.394 |
| Middle Longitudinal Fasciculus | 0.41 | 0.04 | 0.37 | 0.05 | 7.81 | 0.001 | 0.15 | 0.160 |
| Optic Radiation | 0.44 | 0.05 | 0.38 | 0.05 | 12.48 | 0.002 | 0.22 | 0.030 |
| Parieto-Occipital Pontine | 0.51 | 0.05 | 0.45 | 0.05 | 12.34 | 0.002 | 0.22 | 0.030 |
| Striato-Fronto-Orbital | 0.40 | 0.05 | 0.38 | 0.05 | 0.91 | 0.002 | 0.02 | 0.394 |
| Striato-Occipital | 0.46 | 0.06 | 0.39 | 0.05 | 12.24 | 0.002 | 0.21 | 0.030 |
| Striato-Parietal | 0.45 | 0.04 | 0.40 | 0.05 | 11.58 | 0.001 | 0.20 | 0.037 |
| Striato-Post-Central | 0.47 | 0.05 | 0.41 | 0.05 | 6.66 | 0.002 | 0.13 | 0.211 |
| Striato-Pre-Central | 0.50 | 0.06 | 0.44 | 0.05 | 5.92 | 0.003 | 0.12 | 0.246 |
| Striato-Prefrontal | 0.41 | 0.05 | 0.37 | 0.05 | 3.23 | 0.002 | 0.07 | 0.394 |
| Striato-Premotor | 0.42 | 0.06 | 0.38 | 0.05 | 3.15 | 0.002 | 0.07 | 0.394 |
| Superior Cerebellar Peduncle | 0.49 | 0.05 | 0.41 | 0.04 | 20.00 | 0.002 | 0.31 | 0.002 |
| Superior Longitudinal Fasciculus I | 0.38 | 0.04 | 0.34 | 0.05 | 4.58 | 0.002 | 0.09 | 0.302 |
| Superior Longitudinal Fasciculus II | 0.38 | 0.05 | 0.34 | 0.06 | 6.22 | 0.002 | 0.12 | 0.241 |
| Superior Longitudinal Fasciculus III | 0.39 | 0.05 | 0.34 | 0.05 | 6.82 | 0.002 | 0.13 | 0.207 |
| Superior Thalamic Radiation | 0.57 | 0.07 | 0.50 | 0.06 | 7.64 | 0.004 | 0.15 | 0.165 |
| Thalamo-Occipital | 0.44 | 0.05 | 0.37 | 0.05 | 12.49 | 0.002 | 0.22 | 0.030 |
| Thalamo-Parietal | 0.45 | 0.04 | 0.39 | 0.05 | 11.68 | 0.001 | 0.21 | 0.036 |
| Thalamo-Post-Central | 0.52 | 0.06 | 0.45 | 0.05 | 8.32 | 0.003 | 0.16 | 0.132 |
| Thalamo-Pre-Central | 0.52 | 0.06 | 0.46 | 0.06 | 6.95 | 0.003 | 0.13 | 0.206 |
| Thalamo-Prefrontal | 0.41 | 0.05 | 0.37 | 0.05 | 4.36 | 0.002 | 0.09 | 0.302 |
| Thalamo-Premotor | 0.42 | 0.05 | 0.38 | 0.05 | 5.18 | 0.002 | 0.10 | 0.277 |
| Uncinate Fasciculus | 0.43 | 0.05 | 0.39 | 0.05 | 3.22 | 0.002 | 0.07 | 0.394 |

Mean FDC values per tract, collapsing across hemisphere for tracts automatically delineated separately for left and right hemisphere by TractSeg. Univariate ANCOVAs involved age at scanning and sex as between-subjects covariates of no interest; **key:** LGI-1 aLE: anti-leucine-rich glioma inactivated autoimmune limbic encephalitis; CTR: healthy controls; FDC: Fiber Density and Cross-section; p-corr: Bonferroni-Holm correction for multiple testing.
